# Ultra-Rapid and Specific Gelation of Collagen Molecules for Transparent and Tough Gels by Transition Metal Complexation

**DOI:** 10.1101/2023.03.21.533590

**Authors:** Tomoyuki Suezawa, Naoko Sasaki, Nazgul Assan, Yuta Uetake, Kunishige Onuma, Hidehiro Sakurai, Ryohei Katayama, Masahiro Inoue, Michiya Matsusaki

## Abstract

Collagen is one of the main components of tumor stromal tissues with a high elastic modulus, but there have been limitations when attempting to fabricate a tough collagen gel with cells like a cancer stroma. Here, we demonstrate the rapid and specific formation of collagen gels with high transparency and high elastic modulus by transition metal complexation within minutes. Transition metal ions such as K_2_PtCl_4_ exhibited rapid gelation due to the formation of a cross-linked network of the collagen triple-helix by Pt– O and/or Pt–N bonds. Interestingly, type I to IV collagens showed rapid gelation, while other extracellular matrices and DNA did not exhibit this phenomenon, suggesting the importance of intermolecular interaction in a rigid triple-helix structure. Live imaging of colon cancer organoids in three-dimensional culture indicated a collective migration property with modulating high elastic modulus, suggesting activation for metastasis progress. This technology that facilitates deep-live observation and mechanical stiffness adjustment will be useful as a new class of scaffolds.

**Teaser:** Transparent collagen gels with tunable mechanical properties allow deep-live observation of cells cultured in a tough environment like our bodies.

## Introduction

Extracellular matrices (ECMs) are key molecules for regulating the cell microenvironment, controlling cell functions such as adhesion, migration, growth, differentiation, cell-cell interaction, protein secretion, and metabolism (*1, 2*). Collagen (Col) is an important fibrillar ECM as it is a tension-resisting element which maintains tight and stable tissue structures (*3*). Col is a main ECM component of connective (stromal) tissues including tumors (*4*). Recently, increase in ECM stiffness caused by increasing the secreted amount of Col has been found to be one of the strongest risk factors for cancer progression (*5, 6*). For example, although normal breast has a stiffness of only 0.8 kPa, nodal metastasis rates of 396 breast cancer patients ranged from 7 % for tumors with a mean stiffness of < 50 kPa to 41 % for tumors with a mean stiffness of > 150 kPa (*7*). Moreover, increased the elastic modulus of colorectal cancer tissues of 106 patients showed a strong correlation with T stage, denoting the size and extent of the main tumor, and the median values of the elastic moduli at T1 and T4 stages were 2.81 (max. 3.96) and 13.8 (max. 68.0) kPa, respectively (*8*). The reason for these stiffness increases depending on cancer progression has been clarified as increasing Col secretion (*5, 6*). Accordingly, cytocompatible control of the elastic modulus of Col hydrogels for the three-dimensional (3D) culture of cancer cells has been keenly sought after.

To prepare Col hydrogels, covalent bond formation has been generally formed by using a chemical crosslinker such as glutaraldehyde (GA) (*9*) or 1-ethyl-1-3-(3-dimethylaminopropyl) carbodiimide hydrochloride (EDC)-*N*-hydroxysuccinimide (NHS) (*10*), an enzyme such as transglutaminase (*11*) or peroxidase (12), and UV irradiation (13). However, the elastic modulus of the obtained gels is not high and the crosslinkers show cytotoxicity. Genipin has been reported as a cytocompatible crosslinker (*14*), but the obtained gel is stained blue by the reacted genipin derivatives. There have been a few reports on physical crosslinkers, but the elastic moduli of the obtained gels were low (*15*). To avoid the cytotoxicity issue of crosslinkers, a physical crosslinking method by neutralization and heating of a collagen acidic solution has been used as a standard method of 3D-cell culture. However, tough Col hydrogels with over tens kPa modulus have generally not been constructed because of the low dissolving property of Col molecules into an aqueous solution. Col can be dissolved in an acidic solution with concentrations of 5–10 mg/ml at most and subsequent neutralization induces gelation. Accordingly, Col hydrogels below 1.0 wt% concentration are commonly used for 3D culture of cancer cells so their elastic modulus is lower than 1.0 kPa (*16*). We recently reported a unique approach, “sedimentary culture”, using a collagen microfiber (CMF) to fabricate large-scale engineered tissues (*17-19*). The millimeter-sized tissues with high ECM density were easily obtained by centrifugation of cells and CMFs, but the maximum elastic modulus of the obtained tissues was around 10 kPa, much lower than that of cancer at the late stage.

Another limitation of Col gel culture is observation because, although it is a standard method of 3D-cell culture, the gels cannot be observed clearly at great depth even by confocal laser scanning microscopy (CLSM) due to their opaque color. Although transparent collagen gels have been reported in the field of cornea (*20, 21*), 3D-cell culture in the transparent gels has yet to be achieved. Live observation at great depth is still a major challenge in Col gel culture.

In this study, we demonstrate for the first time the rapid gelation of Col solutions by transition metal complexation. Transition metal ions form a complex between a nitrogen or oxygen atom of inter/intra Col molecules, whereas basic metal ions do not. Although interaction between *cis*-diamminedichloroplatinum (II) (cisplatin), which is a well-known anticancer agent, and a Col model peptide has been reported (*22*), there are no reports on crosslinking of natural Col fibers to rapidly form a gel by complexation with transition metal ions. The elastic modulus of the obtained transparent Col gels is widely controllable by varying the metal ion concentration up to 1.8 MPa. Furthermore, live imaging of a patient-derived colon cancer organoid showed activated morphology of the collective migration at 162 kPa, which is a higher modulus range than that of a colon cancer at the T4 stage. This rapid-gelation technology for modulus-controllable and transparent Col gels will provide a new class of 3D-cell cultures for drug assessment and basic cancer biology.

## Results

### Rapid Gelation of Col-TM gels

Type I collagen nanofibers (CNFs) (*23*) which undergo a reversible sol-gel transition from 4 °C to 37 °C under neutral pH conditions were used in this study to avoid the influence of pH on gelation because conventional type I collagen needs to be dissolved in an acidic solution at 4 °C and then both neutralization and heating are necessary for its gelation. When K_2_Pt(II)Cl_4_ solution was added to the CNF solution under a stirring condition, the solution quickly solidified (within 3 min) and the stirring bar was completely stopped (Fig. 1 and Movie S1), suggesting intermolecular crosslinking of CNFs by K_2_Pt(II)Cl_4_ The obtained gels were completely transparent and stably maintained even after flipping the bottle. The elastic modulus of the obtained gels increased with increasing K_2_Pt(II)Cl_4_ and CNF concentrations and reached a plateau at 0.1 mM, independent of CNF concentration (Fig. 2A). Surprisingly, the elastic moduli evaluated by compression tests exceeded 1.0 MPa even in a wet gel condition and the maximum value was 1.78 MPa in the 1.0 wt% CNF and 1.0 mM K_2_Pt(II)Cl_4_ condition. Even the lowest 0.2 wt% CNF condition yielded 162 kPa at 0.5 mM K_2_Pt(II)Cl_4_, a much higher value than that of the conventional 1.0 wt% Col hydrogels with a 1.0 kPa modulus prepared by neutralization and heating (*16*).

**Fig. 1.**
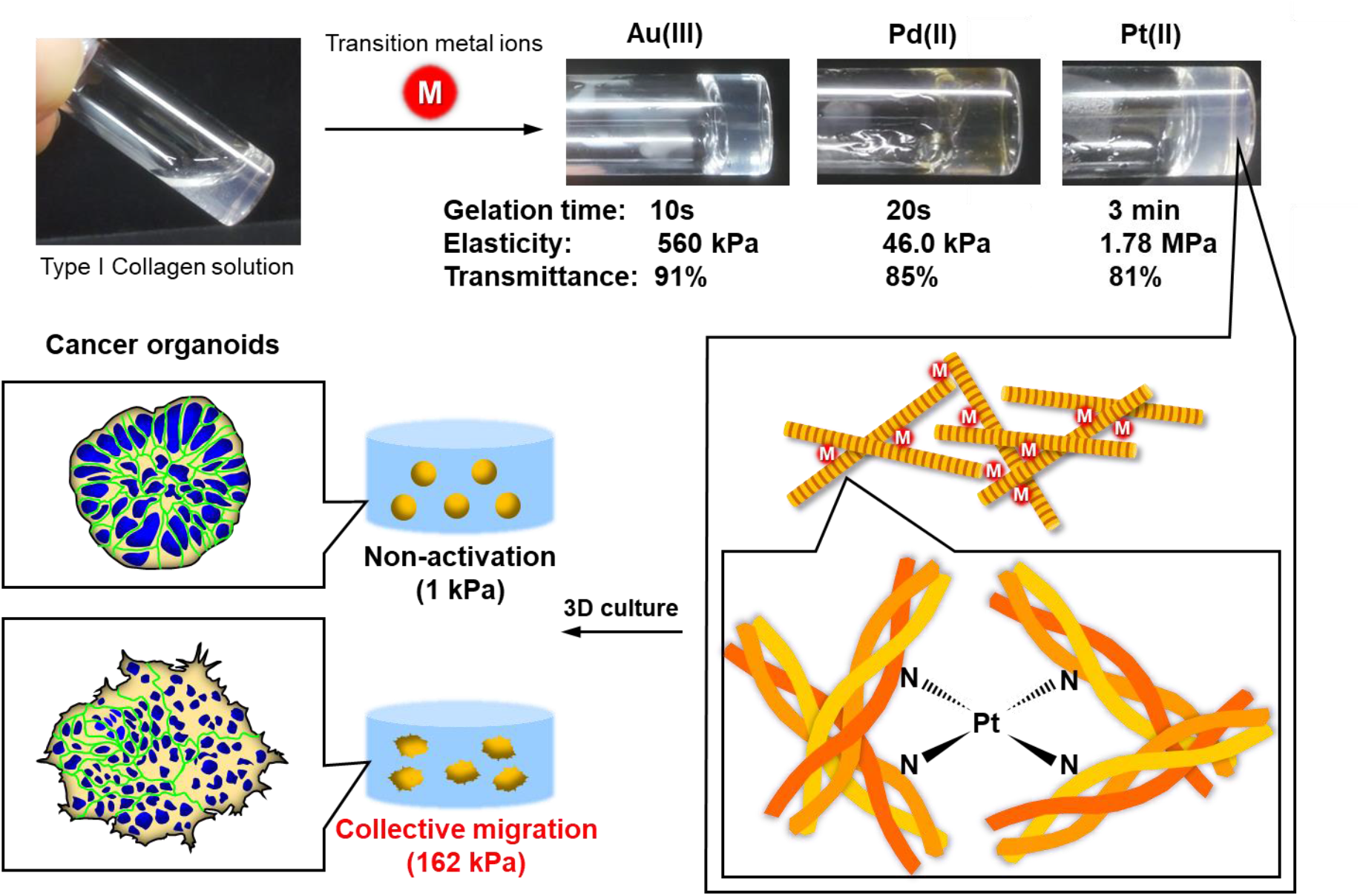
Structural illustration of Col I-TM gels. CNF solution is quickly crosslinked by the coordinate bonding of transition metal ions to nitrogen or oxygen atoms of collagen molecules, resulting in a transparent gel with high elasticity. The modulus-controllable properties of this gel are useful for mechanobiology assays with 3D cultures of cancer organoids.

**Fig. 2.**
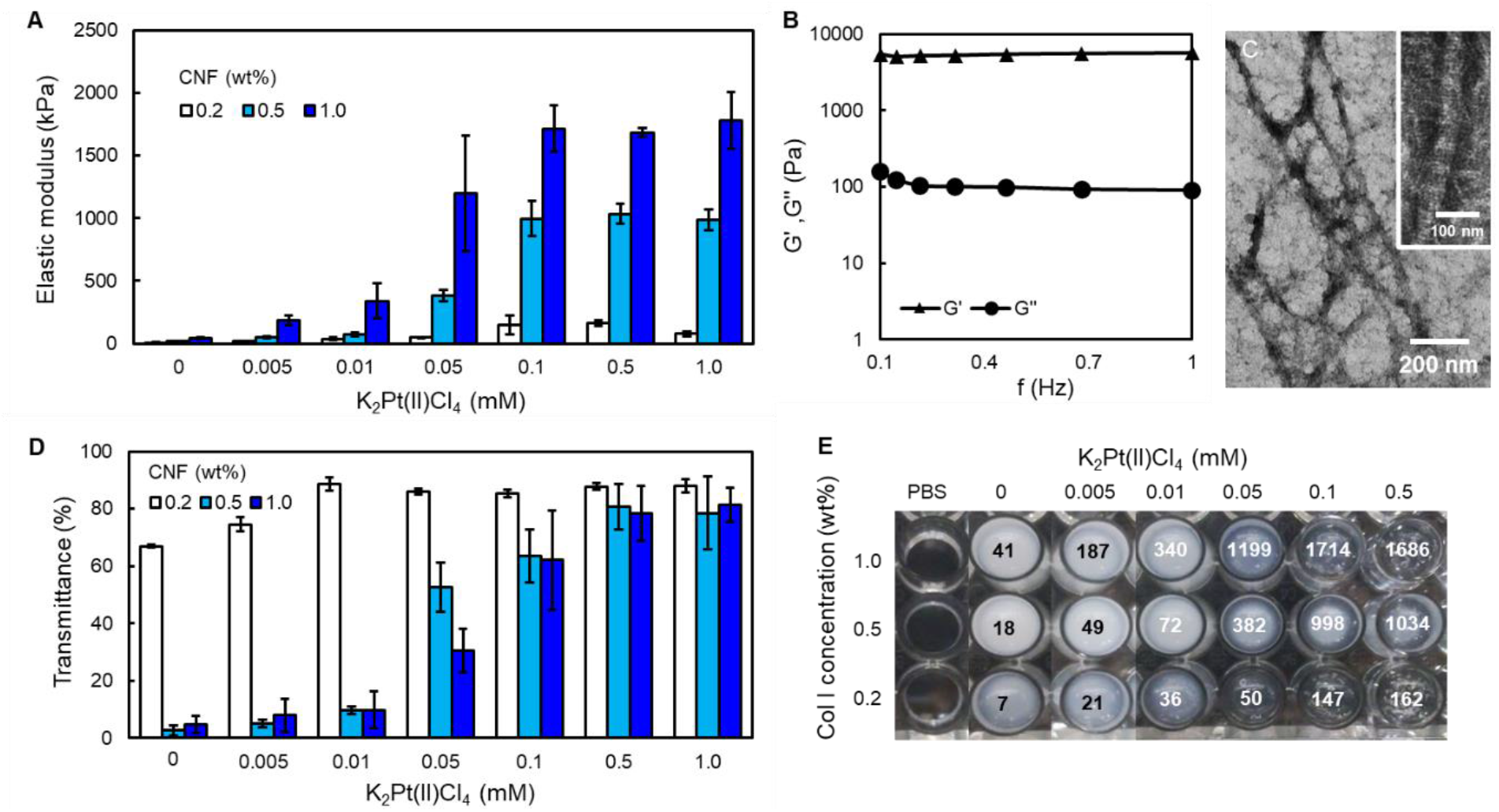
Characterization of Col-Pt gels. (**A**) Elastic moduli of Col-Pt gels constructed under 0.005-1.0 mM K_2_Pt(II)Cl_4_ and 0.2-1.0 wt% CNF conditions at r.t. (n=3). (**B**) Viscoelasticity of Col-Pt gels constructed at 0.5 mM K_2_Pt(II)Cl_4_ and 1.0 wt% CNF at 37°C. (**C**) TEM images of Col-Pt gels constructed at 0.1 mM K_2_Pt(II)Cl_4_ and 0.2 wt% CNF at 37°C. The gels were stained with uranyl acetate. (**D**) Transmittance at 500 nm and (**E**) photograph of the obtained Col-Pt gels in 96 microwell at 37 °C (n=3). The values in each well are elastic moduli (kPa).

Interestingly, almost all water-soluble transition metal ions such as group 10 (Ni(II), Pd(II), and Pt(II)/(IV)), group 11 (Cu(II) and Au(III)), and group 12 (Zn(II)) showed rapid gelation with a high elastic modulus (Movie S2). Conversely, group 8 (Fe(III)), group 9 (Co(II) and Ir(IV)), and part of group 11 (Ag(I) and Au(0)) did not show this phenomenon which was the same as group 2 (alkaline earth metal: Mg(II), Ca(II), and Ba(II)) (Table S1). From these data, at least multivalent transition metal ions were expected to have the rapid gelation property of Col I molecules. The resulting Col-transition metal (TM) gels were named “Col-TM gels”.

The rheological property of the Col-Pt gels is shown in Fig. 2B. Storage modulus (G’) was stably maintained at around 5,500 Pa under frequency variation, suggesting high surface stiffness. Loss moduli (G’’) were 100–150 Pa, around 50–fold lower than the G’ value, and Tan δ (G’’/G’) were 0.016–0.025, indicating a solid-like, typical chemically bonded property (*24*). TEM observation of Col-Pt gels indicated a crosslinking of collagen fibrils that clearly showed D-band periodicity (Fig. 2C).

To gain an understanding of the transparency of the Col-Pt gels, their transmittance was measured at different K_2_Pt(II)Cl_4_ and CNF concentrations (Fig. 2D). The gels prepared with 0.2 wt% CNF solution all showed transparency greater than 67 %. The gel transparency increased with increasing Pt ion concentration, reaching a plateau of 90 % at a Pt ion concentration above 0.01 mM. The gels prepared with higher CNF concentrations also showed the same trend, reaching a plateau of 80 % at a Pt ion concentration above 0.5 mM even though their elastic moduli were higher than 1 MPa (Fig. 2E). The above findings suggest the formation of a homogeneous network inside the gels (*25*).

### Structural analysis of Col-TM gels

In general, Col gels composed of triple-helix molecules have an opaque color due to phase-separation, while gelatin gels composed of dissociated single collagen molecules with a random structure have transparent properties. To confirm the triple-helix structures of the Col-TM gels, circular dichroism (CD) spectra of the gels using HAu(III)Cl_4_ were analyzed under different metal ion concentrations (Fig. S1). The CNF exhibited a typical positive peak at 225 nm corresponding to a triple-helix structure, and the peak intensity showed no change with increasing TM ion concentration, suggesting that the triple-helix structure was fully maintained by this gelation. Since the Col-Pt gels prepared by K_2_Pt(II)Cl_4_ also indicated the same triple-helix CD spectra (Fig. 3A), the crosslinking by TM ions did not affect the structure of the collagen molecules.

**Fig. 3.**
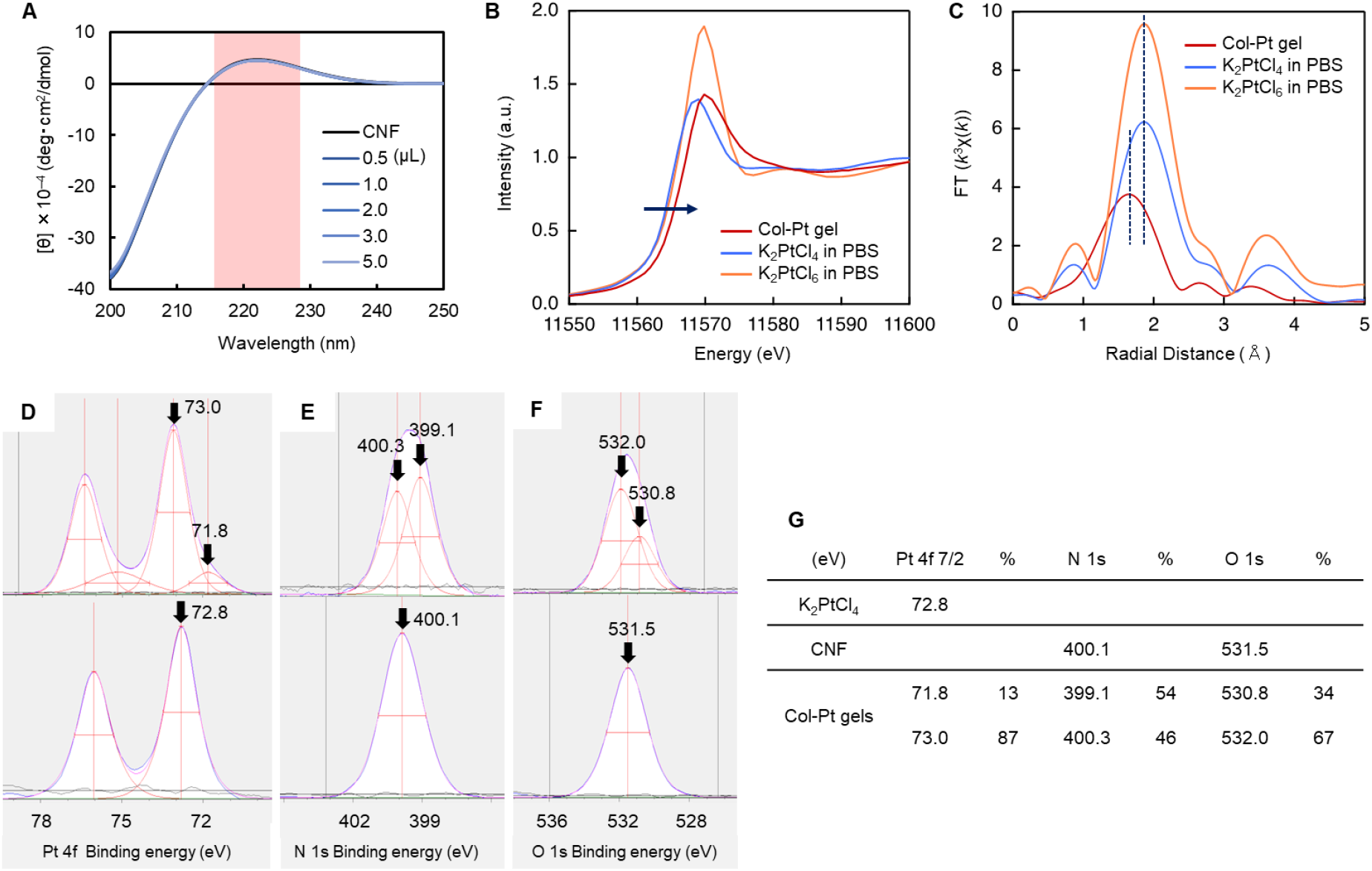
Structural analysis of Col I-Pt gels. (**A**) CD spectra of a CNF solution within different concentrations of Pt(II) at room temperature. (**B**) XANES and (**C**) EXAFS of K_2_PtCl_4_ and K_2_PtCl_6_ in PBS and Col-Pt gels. XPS spectra of (**D**) Pt 4f_7/2_ of Col-Pt gel (top) and K_2_Pt(II)Cl_4_ powder (bottom), (**E**) N 1s of Col-Pt gels (top) and CNF gels (bottom) and (**F**) O 1s of Col-Pt gels (top) and CNF gels (bottom), respectively. The Col-Pt gels were prepared by 50 mM Pt(II) ions and 0.2 wt% CNF. CNF gels were prepared using 0.2 wt% CNF solution by incubation at 37 °C overnight. (**G**) Binding energy peak shift of (**D**) Pt 4f_7/2_, (**E**) N 1s and (**F**) O 1s of Pt(II) ions, CNF, and Col-Pt gels prepared by 50 mM Pt(II) ions and 0.2 wt% CNF.

To understand the role of Col gelation, other commercially available types of collagen molecules such as type II, III, and IV from different animal species were mixed with K_2_Pt(II)Cl_4_ in the same way as for Col I (Table S2). All collagen molecules in phosphate buffered saline (PBS) at pH 7.4 quickly gelled with the same transparency as the CNF solution, suggesting the importance of the triple-helix structure for Pt crosslinking. Furthermore, other types of ECMs, collagen model peptides, polysaccharides, synthetic polymers, and DNA were selected to evaluate their potential for gelation by Pt ions (Table S3). Although gelatin is a component of the single collagen α chain in the triple helix, it did not show gelation, suggesting the importance of triple-helix molecules (Movie S3). A collagen model peptide ((POG)_10_) had a collagen triple-helix structure with a positive peak at 225 nm in the CD spectra but did not show gelation probably due to its much lower molecular weight. DNA has a double-helix structure but yielded the same results. Laminin and fibronectin are well known cell adhesive ECMs with the same high molecular weight as collagen, however gelation was not observed. Natural polysaccharides, heparin and alginate, and the synthetic polymer polyacrylate were also used because of their abundance of oxygen atoms that can be expected to form coordinate bonds with Pt ions. However, none of the molecules exhibited gelation in the same manner as collagen molecules. These results strongly suggest that this method reveals high molecular selectivity of the collagen triple helix. The difference between DNA and the collagen triple helix is interesting because both have a rigid helical structure and DNA is well known for its ability to make coordinate bonds with cisplatin Pt(II) (*26*). Since the triple helix of collagen is a linear and more packable molecule than DNA, tight packing of the huge molecules is considered to be of important.

Because gelatin molecules dissolve well in water and partly form a triple-helix structure at the edge of the molecules in a high concentration condition, higher concentrations of both gelatin and K_2_Pt(II)Cl_4_ were used as shown in Table S4. In particular, gelatin showed collagen-like gelation at over 10 wt% gelatin and 10 mM Pt ions, likely due to the high packing of the partially formed triple-helix structures. To understand molecular entanglement, the viscoelasticity of both Col I and gelatin were measured at different concentrations (Table S5). Interestingly, Col I showed around 20-fold higher viscoelasticity than gelatin, probably due to intermolecular hydrogen bond formation of huge linear collagen nanofibrils (*3*). These results clearly suggest the importance of tight packing huge collagen fibrils for TM ion crosslinking.

To confirm the local geometry around platinum atom in Col-Pt gels, Pt L_3_-edge X-ray absorption spectroscopy (XAS) was performed. Col-Pt gel prepared from K_2_Pt(II)Cl_4_ was incubated for 1 week at 4 °C to form the tough gel, and XAS experiments were conducted with fluorescent method. Focusing on the X-ray absorption near edge structure (XANES) region, the absorption edge of the Col-Pt gel appeared at 11565 eV, which is applox. 1 eV higher than that of K_2_Pt(II)Cl_4_ in PBS (Fig. 3B). Since the first peak of Pt L_3_-edge XAS corresponds the electric dipolar transition from filled 2p_3/2_ to unfilled 5d oritals, the observed chemcal shift denotes the increase of 5d energy level of platinum. Therefore, this result suggested the ligand exchange from the chloride atoms to electron-donating elements, such as oxygen or nitrogen included in collagen structure, occured during the gelation process. In addition, the peak intensity of Col-Pt gel was almost same as that of K_2_Pt(II)Cl_4_, and lower than that of K_2_Pt(IV)Cl_6_, indicating that the platium atom in Col-Pt gel possesses a square-planer geometry which is typical for Pt(II) complexes. The extended X-ray absorption fine strucuture (EXAFS) of Col-Pt gel, K_2_Pt(II)Cl_4_, and K_2_Pt(IV)Cl_6_ were shown in Fig. 3C. In the case of K_2_Pt(II)Cl_4_ and K_2_Pt(IV)Cl_6_, a peak was observed at 1.9 Å which corresponds the Pt–Cl scattering. Meanwhile, the peak was shifted to 1.6 Å after gel formation. This result indicates that the nearest elements of platinum were replaced by light elements, such as oxygen and nitrogen, showing a good agreement with the result of XANES data. To sum up, the platinum in the Col-Pt gels has a square planar structure and is found to bind to nitrogen and oxygen atoms present in collagen. Hence, it is thought that the platinum plays a role in connecting the collagen fibers having triple-helix structure to form a cross-linked network as shown in Figure 1, that is considered to be an origin of the toughness of the thus-fabricated Col-Pt hydrogels. Noted that the XANES spectra of Pt(II) in Col-Pt gels after 1 h and 1 week of incubation were almost the same, indicating that 1 h incubation was sufficient for ligand exchange (Fig. S2).

To confirm coordinate bond formation between Pt(II) and N or O atoms in collagen molecules, the binding energy was observed by X-ray photoelectron spectroscopy (XPS) (Figs. 3D-G). In this technique, the 4f orbitals of Pt are split by the interaction of orbital magnetic moment and spin, and the lower energy peaks (4f_7/2_) are used to evaluate the binding energy (*27*). Although the peak for K_2_Pt(II)Cl_4_ powder was a single peak at 72.8 eV which was the same as in a previous report (*27*), two peaks at 71.8 and 73.0 eV were estimated by curve fitting, suggesting the coordinate exchange from Cl atoms to the other atoms (Fig. 3D). The peak shift to lower energy denotes a decrease in electrical binding from the nucleus (the reduction of Pt), suggesting coordinate exchange from Cl to N or S (higher to lower electronegativity) (*28*). On the other hand, the peak shift to higher energy denotes a coordinate exchange from Cl to O (lower to higher electronegativity). The peak splitting was also observed on N 1s binding energy in CNF gels (Fig. 3E). The peak shift from 400.1 eV to 399.1 eV denotes the reduction of N atoms, suggesting the coordinate bond formation with Pt atoms. This reduction was also found on O 1s binding energy in CNF gels (Fig. 3F). The results of XPS spectra summarized in Fig. 3G clearly indicated the coordinate bond formation between Pt(II) and N and O atoms in collagen molecules.

To measure the amount of Pt(II) ions used for crosslinking to form the Col-TM gels, inductively coupled plasma (ICP) spectroscopy was conducted (Fig. S3). The amount of Pt(II) ions inside the gels increased with increasing CNF concentration, while excess Pt(II) ions in washing buffers decreased, suggesting an increase in the crosslinking bonds inside the gels by Pt(II)-collagen coordinate bonds (Fig. S3A). In the case of increasing Pt(II) ion concentrations, the Pt(II) ions inside the obtained gels gradually increased and then reached a plateau at concentrations exceeding 0.5 mM Pt(II) ions (Fig. S3B). The theoretical crosslinking bond number in the Col-TM gels prepared by 0.5 wt% CNF and 0.5 mM K_2_Pt(II)Cl_4_ was calculated as 2.4 × 10^−7^ by the 12 μg of Pt(II) ions because Pt(II) ions form tetragonal coordinate bonds. Amino acids that could possibly be crosslinked in collagen molecules by Pt(II) ions are expected to be hydroxyproline, serine, tyrosine, threonine, histidine, lysine, hydroxylysine, asparagine, aspartic acid, glutamine, glutamic acid, methionine, and cysteine based on the results of XPS spectra indicating Pt–N and Pt– O formation (Fig. 3D–G) and previous reports on Pt–S formation (*28*). Since the ratio of these amino acid residues in collagen molecules was estimated at 321:1,000 (*29, 30*), the theoretical maximum crosslinking bond in the gels is estimated to be 5.1 × 10^−6^, around 21-fold higher than the above estimated number, probably due to the steric hindrance of huge linear collagen fibrils and segregation of the possible amino acids to the fiber interior.

### Healing property of Col-TM gels

For the application of the Col-TM gels, the self-healing property at the cleavage surface was expected because of the noncovalent nature of the coordinate bond as previously reported (*31*). Unfortunately, however, the cleavage surfaces of the Col-TM gels did not attach well even after overnight incubation. We therefore added additional solution of PBS, 1.0 mM K_2_Pt(II)Cl_4_ and 1.0 wt% CNF at the cleavage interfaces and the gels were then incubated at r.t. for 1 h (Fig. 4A). In the case of PBS and K_2_Pt(II)Cl_4_, the adhesive phenomon was not observed at all. However, in the solution with CNF, adhesion of the cleavage surfaces was clearly observed within 10 min, which might have been due to excess Pt(II) ions, as shown in Fig. S3. Additional Col molecules in between the cleavage surfaces were crosslinked by the excess Pt(II) ions, because the crosslinkable amino acid residues in the Col molecules seemed to be already wholly crosslinked during the Col-TM gel formation.

**Fig. 4.**
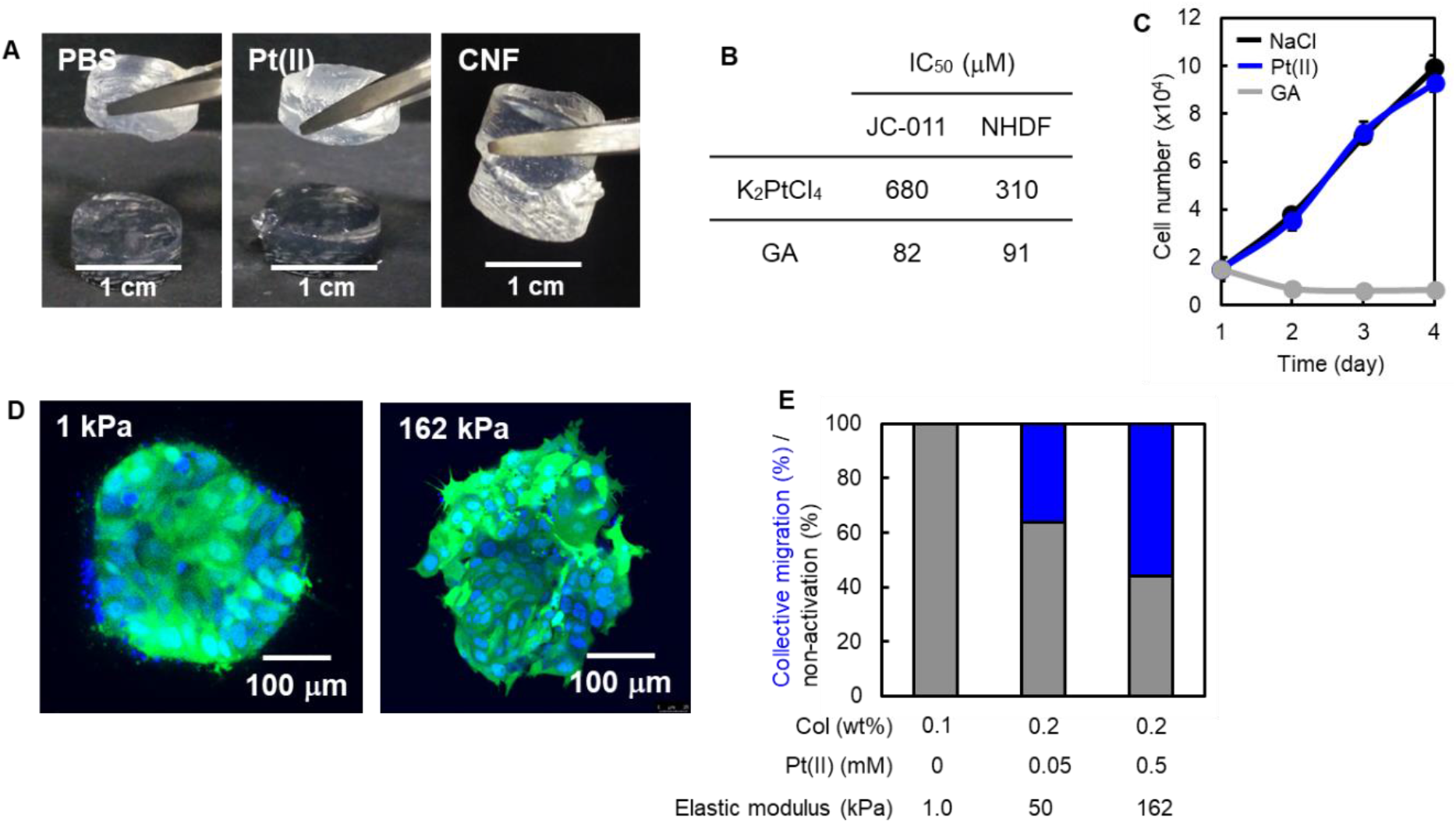
Application of Col-TM gels. (**A**) Healing property on the cleavage surfaces of Col-Pt gels constructed with 1.0 wt% CNF and 1.0 mM Pt(II) by adding a further1.0 wt% CNF. (**B**) IC_50_ values of K_2_Pt(II)Cl_4_ and GA against a patient-derived colon cancer cell line (JC-011) and NHDFs in DMEM containing 10 % FBS with varied concentrations after 3 days of culture at 37 °C (n=3). The values were calculated from Fig. S4. (**C**) Cell proliferation profiles of NHDFs in DMEM containing 10 % FBS with 0.1 mM NaCl, GA, or K_2_Pt(II)Cl_4_ for 4 days of culture (n=3). (**D**) CLSM live imaging of Hoechst staining for 3D cultured EGFP-labeled organoids in stiff (162 kPa) Col-Pt gels constructed by 0.2 wt% CNF with 0.5 mM K_2_PtCl_4_ solutions. The control soft gels (1 kPa) without Pt (II) ions were constructed by a commercial 0.1 wt% type I collagen solution by neutralization and heating at 37 °C. (**E**) Mean percentage of both collective migration and non-activation of the organoids after 3D culture inside the gels constructed by varied Col and Pt(II) concentrations (n=5∼9).

### Cytocompatibility of transition metal ions

Finally, the cytocompatibility of the Col-TM gels was evaluated to understand their potential application in cancer cell culture. Patient-derived colon cancer cell line (JC-011) and normal human dermal fibroblasts (NHDFs) were cultured in dulbecco’s modified eagle medium (DMEM) containing 10% fetal bovine serum (FBS) and K_2_Pt(II)Cl_4_ or glutal aldehyde (GA) at varied concentration for 3 days, and then half maximal inhibiory concentration (IC_50_) of Pt(II) ions and GA was estimated (Figs. 4B and S4). GA was used as a control of typical chemical crosslinker. IC_50_ values of Pt(II) ions to JC-011 and NHDFs were 680 and 310 μM respectively and these were 8.2 and 3.4-fold lower cytotoxicities than those of GA, suggesting the potential of the Col-TM gels for 3D cell cultures at high elastic modulus. To understand the effect of Pt(II) ions to cell growth, cell proliferation profiles of NHDFs were evaluated for 4 days in DMEM containing 10% FBS with 0.1 mM NaCl, K_2_Pt(II)Cl_4_ or GA, respectively (Fig. 4C). Pt(II) ions revealed exactly the same growth profile as NaCl, but GA caused a decrease in cell number, suggesting strong cytotoxicity. These data clearly demonstrate that Pt(II) ions have high pontetial for 3D-cancer cell cultures with controllable elastic modulus.

### 3D culture of cancer organoids in Col-TM gels

To understand the stiffness effect of the collagen-gel matrix to the phenotype of cancer cells, patient-derived colorectal cancer (CRC) organoids were sandwich-cultured by the Col-TM gels at a stiff (162 kPa) elastic modulus for 7 days and observed by CLSM in live conditions (Fig. 4D). We previously reported a cancer tissue-originated spheroid (CTOS) method, in which cell-cell contact of cancer cells were maintained throughout the preparation process (*32*). The organoids prepared by CTOS method is useful for drug screening and personalized therapy because it retains the properties of cancer cells in the patient’s original tumor (*33, 34*). When CRC organoids were cultured in control soft collagen gels at 1 kPa, no morphological changes were observed and the cancer cells remained uniformly aligned. However, the 162 kPa Col-TM gels clearly indicated a marked spike and cell migration toward the outer boundary, suggesting morphology of collective migration that is the process by which a group of cells moves together, without completely disrupting their cell-cell contacts. To understand the stiffness effect in detail, the concentration of CNF and K_2_Pt(II)Cl_4_ solutions was varied for 3D culture of the organoids and the ratio of collective migration was estimated from the organoid morphology (Figs. 4E, S5 and S6). The concentration of K_2_Pt(II)Cl_4_ solutions was adjusted to less than 1.0 mM to avoid cytotoxicity. When concentration of K_2_Pt(II)Cl_4_ solutions was increased, the ratio of the collective migration clearly increased regardless of Col concentration. This suggests that collective migration may be induced by the stiffening of the ECM scaffolds in 3D culture. The collective migration is important during morphogenesis, and in pathological processes such as wound healing and cancer cell invasion (*35*). While the molecular mechanisms of collective migration are becoming clear, analysis of the underlying mechanisms of collective migration in cancer is still at an early stage. The importance of mechanical cues in the collective migration has recently been studied, and *in vivo* experiments have revealed that tissue stiffening initiates the epithelial-to-mesenchymal transition that triggers collective migration (*36*). Although biomechanical methods such as 2D culture, micro-patterning, 3D-tube migration, and computational simulation have been reported extensively (*37*), there are no reports of collective migration initiated by stiffening the ECM scaffolds in 3D culture. The 3D culture in the Col-TM gels with controllable environmental ECM stiffness has the potential to accelerate *in vitro* analysis of the underlying mechanism of collective migration in cancer.

## Discussion

We reported the rapid gelation of Col solutions by transition metal complexation with nitrogen or oxygen atoms of inter/intra Col molecules. It has been reported that chrome treatment improves the heat resistance of tanned leather, and the complexation of collagen fibrils and chromium was proposed as the mechanism, but the details remained unclear (*38*). This study reveals the historical mechanism of complexation of transition metal ions with collagen fibers and shows a new application of collagen transparent gelation in the biomedical field. This stiffness-controllable, rapid, and transparent gelation technique may be useful for a new class of biomaterials for tissue engineering and basic biology.

## Materials and Methods

### Preparation of CNF

Pepsin-treated Type I collagen (Col I) sponge were kindly donated by NH Foods Ltd (Osaka, Japan). The 50 mg of type I collagen sponge and 5 ml of PBS (D5652-10L, Sigma–Aldrich, St. Louis, USA) were added into a 15 ml centrifuge tube (Corning, 430791, NY, USA). A φ8 mm blade was set in a homogenizer (AS ONE, VH-10 1-8471-31, Osaka, Japan) and homogenized collagen sponge at a speed of 6 (30,000 rpm) for 6 min. After homogenization, the collagen suspension was incubated at 4 °C for 1 day to dissolve collagen microfibers. After incubation, the solution was centrifuged (5922, KUBOTA, Tokyo, Japan) at 25 °C and 10000 rpm for 3 min to remove bubbles and CNF solution. Other concentrations were prepared in the same manner by changing the amount of collagen sponge or adjusting the concentration after resuspension.

### Gelation of CNF solution by transition metal ions

Five hundred µl of 0.5 wt% CNF solution and 20 µl of 12.5 mM potassium tetrachloroplatinum (II) acid (K_2_PtCl_4_: 206075-1G, Sigma-Aldrich, St. Louis, USA) in PBS to the sample tube (0201-01 Maruemu, Osaka, Japan). Then, they were mixed using a pipetter (SA05825, Gilson, Middleton, USA) for high viscous solution, 0.5 mM Pt(II)-0.5 wt% CNF gel was prepared at 4 °C for 1 day. For the following metal species, gels were prepared in the same manner and the concentration of each metal was adjusted by the amount of metal solution. Magnesium chloride (MgCl_2_: 136-03995, WAKO, Osaka, Japan), Calcium chloride (CaCl_2_: 039-00475, WAKO, Osaka, Japan), Barium (II) chloride (BaCl_2_: 026-00185, WAKO, Osaka, Japan), Iron (III) chloride (FeCl_3_: 010-40684, KISHIDA CHEMICAL, Osaka, Japan), Cobalt (II) chloride (CoCl_2_: 035-10982, WAKO, Osaka, Japan), Iridium (IV) chloride (IrCl_4_: 092-02921, WAKO, Osaka, Japan), Nickel (II) chloride (NiCl_2_: 654507-5G, Sigma-Aldrich, St. Louis, USA), Palladium (II) chloride (PdCl_2_: 7647-10-1, TCI, Tokyo, Japan), Cis-diamine dichloroplatinum (II) (Pt(NH_2_)Cl_2_: 039-20093, WAKO, Osaka, Japan), Potassium hexachloroplatinum (IV) (K_2_PtCl_6_: 206067-1G, Sigma-Aldrich, St. Louis, USA), Copper (II) chloride (CuCl_2_: 032-04142, WAKO, Osaka, Japan), Silver (I) nitrate (AgNO_3_: 000-70555, KISHIDA CHEMICAL, Osaka, Japan), Gold nanoparticles (Au NP: 752568, Sigma–Aldrich, St. Louis, USA), Gold (III) acid tetrahydrate (HAuCl_4_: 077-00931, WAKO, Osaka, Japan), and Zinc (II) chloride (ZnCl_2_: 261-00272, WAKO, Osaka, Japan).

Gelation was visually confirmed from naked-eye observation after tilting the bottles. The classification of gelation condition is as follows: the circle means complete gelation, the triangle means partly gelation, and the x means non-gelation. The CNF gels by each metal species were prepared in 24 well inserts (Corning, clear-3470, NY, USA) and then the elastic moduli of the obtained gels were measured after 1 day of incubation at 37 °C.

## Supplementary Materials

**This PDF file includes:**

Materials and methods

Effect of TM species on gelation

CD spectrum of Col-Au gels

Effect of collagen species and type on gelation

Effect of polymer type on gelation

Effect of gelatin concentration on gelation

Viscoelasticity of CNF and gelatin solutions

Pt L_3_-edge X-ray absorption spectroscopy of Col-TM gels

ICP data of Col-Pt gels

Cell viability assay

3D culture of organoids in Col-Pt gels

Gelation movies of Col-TM gels

## Acknowledgments

Colorectal cancer surgical specimens were collected from patients who underwent surgery of primary and metastasized colorectal tumor. The patients submitted written informed consent for genetic and biological analyses, which were performed in accordance with the protocols approved by the institutional review board (IRB) of Japanese Foundation for Cancer Research (#2013-1093).

XAS measurements were performed at the BL-9A of KEK under the approval of the Photon Factory Program Advisory Committee (proposal no. 2020G006).

## Author contributions

T.S., N.S., N.A., Y.U. and H.S. contributed to Col-TM gel formation and characterization. K.O., R.K., and M.I. contributed to cytocompatibility and cancer organoid culture. M.M. wrote the manuscript and all authors reviewed and edited the manuscript.

## Data and materials availability

All data are available in the main text or the supplementary materials.

## Supplementary Materials for

## S1. Materials and Methods

### Gelation of polymers and proteins solution by transition metal ions

To evaluate the gelation properties of the following polymers and proteins, the same gelation procedure using 0.5 wt% polymer in phosphate buffer solution (PBS, pH=7.4) and 0.5 mM K_2_PtCl_4_ was performed. Pig skin type III collagen (Col III: KP-5005, Nitta Gelatin, Osaka, Japan), Pepsin-untreated porcine skin type I tropocollagen (Tropo Col I: donated by NH Foods Ltd), Human Col I (donated by NH Foods Ltd), Human placental type IV collagen (Col IV: C7521, Sigma–Aldrich, St. Louis, USA), Chicken cartilage type II collagen (Col II: donated by NH Foods Ltd), Bovine dermis Col I (ASC-1-100-100PW, Nippi, Tokyo, Japan), collagen mimic peptide: (POG)_10_ (4033, Peptide Institute INC., Osaka, Japan), Laminin (354259, Corning, NY, USA), Fibronectin (F2006-5MG, Sigma-Aldrich, St. Louis, USA), Heparin (081-00136, WAKO, Osaka, Japan), Alginic acid (180947-100G, Sigma-Aldrich, St. Louis, USA), Polyacrylic acid (169-18591, WAKO, Osaka, Japan), DNA from salmon sperm (043-31381, WAKO, Osaka, Japan), and Gelatin (077-03155, WAKO, Osaka, Japan) were used in this study.

Gelation of 5.0–20 wt% gelatin solution was also evaluated by K_2_PtCl_4_ at 1–50 mM. In addition, the viscosities of 0.2–1.0 wt% CNF and 5.0–20 wt% gelatin solutions were measured using a Rheometer (HAKKE RheoStress 6000, Thermo Scientific).

### Circular Dichroism (CD) Spectroscopy

The CD spectra were acquired on a spectrometer (J-725, JASCO, Tokyo, Japan) and using a quartz cuvette with 10 mm path length. The 0.5–5.0 µl of 12.5 mM K_2_PtCl_4_ and HAuCl_4_ solutions were added in 3 mL of 0.005 wt% CNF in PBS for CD spectrum measurement. Mean residue ellipticity (MRE, deg·cm^2^·dmol^−1^) values were calculated using the following equation, where *θ* is the ellipticity (deg), *l* is the path length (m), *C* is the collagen and gelatin concentration (M), *N* is the number of residues.

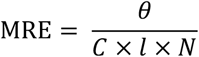

### X-ray Absorption Spectroscopy (XAS)

Pt L_3_-edge X-ray absorption spectroscopy (XAS) experiments were performed at the BL-9A of KEK using Si(111) double-crystal monochromatized synchrotron radiation. All XAS experiments were carried out using fluorescent mode at room temperature. The ionization chamber was used to measure the intensity of the incident X-ray. The X-ray fluorescence was monitored with a 7-element silicon drift detector (SDD). XAS analyses were conducted using Demeter package, a comprehensive system for processing and analyzing X-ray absorption spectroscopy data (*S1-S3*). Background removal and normalization of raw data were performed by cubic spline method using Athena software. *E*_0_ was defined as photon energy at the absorption edge where μT = 0.5 in the normalized μ(*E*) spectrum. The *E*_0_ value of the Pt foil was set to 11564 eV for photon energy calibration. The extracted *k*^3^-weighted EXAFS oscillation was Fourier transformed in the *k*-range 3.0–8.0 Å^−1^.

To a vial (0201-01, Maruemu, Osaka, Japan) containing 1.0 wt% CNF, (POG)_10_, or gelatin in PBS (pH = 7.4, 1.92 mL) was added a PBS containing Pt ion (K_2_PtCl_4_ or K_2_PtCl_6_) (5 mmol/L, 80 μL). After incubating for 60 min or 1 week at 4 °C, the thus-prepared hydrogel or solution was transfer to a plastic bag (polypropylene), and subjected to XAS measurement.

### X-ray photoelectron spectroscopy (XPS)

The Pt(II)-CNF dried gel films prepared by K_2_PtCl_4_ were used for XPS measurements. 0.2 M Pt(II)/PBS solution was prepared and then mixed with 0.2 wt% CNF solution in a 1.5 ml tube (3810X, Eppendolf, Hamburg, Germany) to prepare 50 mM Pt(II)-0.2 wt% CNF solution. Before gelation, 50 µL of mixed solution was rapidly dropped onto a cover glass (C218181, MATSUNAMI, Osaka, Japan) and then dried at 37 °C for overnight. For standard of XPS measurements, 50 µl of Au-NPs (752568, Sigma–Aldrich, St. Louis, USA) in PBS was dropped onto all dried samples and dried at 37 °C for 3 hours. The films were fixed onto the XPS stage with a conductive double-sided carbon tape and then measured on a X-ray photoelectron spectrometer (JPS-9010, JEOL, Tokyo, Japan) using X-ray source of Mg.

### Inductively coupled plasma (ICP)

The CNF gels were prepared in 0.1–1.0 mM Pt(II)-0.5 wt% CNF in PBS and 0.5 mM Pt(II)-0.2-1.0 wt% CNF in PBS by K_2_PtCl_4_ in 24 well inserts. Each gel was immersed in 10 ml of PBS for 1 day at room temperature and then washed with fresh 10 ml of PBS. The total 20 ml of supernatant were collected and lyophilized to obtain as a powder. The obtained powders were completely dissolved in 5 ml of 0.1 M HCl. The obtained Pt(II)-CNF gels were minced and then completely dissolved in 2 ml of 5 M HCl in 24 well plate (3820-024, IWAKI, Shizuoka, Japan). One ml of the obtained solution was transferred to a 15 ml tube and diluted with 4 ml of 0.1 M HCl. The supernatants and dissolved solutions were transferred to glass tubes (TEST15-105NP, IWAKI, Shizuoka, Japan) and then an ICP atomic emission spectrometer (ICPS-7510, SHIMATZU, Kyoto, Japan) was used to determine the amount of Pt(II) ions inside the gels or supernatants using calibration curve of 0–10 ppm Pt(II)/0.1 M HCl solution.

### Compression test

The elastic moduli of the obtained CNF gels were measured using a compression tester (EZ-Test, SHIMADZU, Kyoto, Japan). CNF solutions were incubated at 37 °C for 1 h in 24 well insert after mixing with transition metal solution, and then immediately subjected to compression testing at room temperature. The elastic modulus *E* (kPa) was calculated using the following equation, where *S* is the slope of the stress-strain curve (mN/mm), *H* is the height of the gel (mm) and *C* is the contact area of the testing jig with the gel (mm^2^)

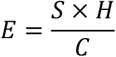

The slope of the graph was selected to follow the rising edge of the stress-strain curve at strains less than 5 %, the height of the gel was measured using a digital caliper (Digimatic Caliper CD-15CP, Mitsutoyo, Kanagawa, Japan), and the contact area of the jig was calculated by integrating the strain values in the stress-strain curve.

### Rheological analysis

The 500 µl of 1.0 wt% CNF gel just after mixing with 0.5 mM K_2_PtCl_4_ was added onto a rheometer and the moduli were measured on a plate geometry with a 20 mm cone at a gap of 0.5 mm using frequency sweep mode in the range of 0.1–1 Hz at 37 °C of stage temperature with 10 Pa stress.

### TEM observation

Col-TM gels were constructed by mixing 0.2 wt% CNF and 0.1 mM K_2_Pt(II)Cl_4_ at r.t. for 10 min. The constructed gels were placed on Cu grids and then dried in vacuum at r.t. for overnight. After drying, the gels were washed with Milli-Q for twice and then the dried samples were stained with 2% uranium acetate. The gels were observed using a transmission electron microscope (H-7000, HITACHI, Tokyo, Japan).

### Transmittance measurement

The 0.005-1.0 mM Pt(II)-0.2-1.0 wt% CNF gels were prepared by K_2_Pt(II)Cl_4_ in 96 well plate (3860-096, IWAKI, Shizuoka, Japan) by incubation at 37 °C for 1 day. To evaluate the transmittance, absorbance at 500 nm was measured using a plate reader and transmittance was calculated using following equation. *A* means the absorbance at 500 nm.

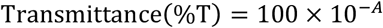

### Patient selection and establishment of patient-derived colon cancer cell line (JC-011)

Colorectal cancer surgical specimens were collected from patients who underwent surgery of primary and metastasized colorectal tumor. The patients submitted written informed consent for genetic and biological analyses, which were performed in accordance with the protocols approved by the institutional review board (IRB) of Japanese Foundation for Cancer Research (#2013-1093). Several pieces of the surgically resected tumors were immediately transferred into the ice-cold culture medium with antibiotic-antimycotic (Gibco). Tumor tissues were cut into small fragments, and enzymatically digested with collagenase/dispase (Roche) and DNase I in StemPro hESC culture medium (Invitrogen) for 60 minutes. After washing with antibiotic–antimycotic and 0.2% BSA-containing PBS, the cell pellets were cultured in the StemPro hESC medium supplemented with 10 μmol/L of Y-27632 to establish the patient-derived JC-011 cell line. Before subsequent experiments, the cells were subcultured in the 1:1 mixed medium of RPMI1640 and Ham’s F-12 supplemented with 10 % fetal bovine serum (FBS), 1 % antibiotic–antimycotic and 10mM Hepes buffer until the coexisting stromal cells were scarcely detected under the microscope.

### Cytotoxicity assay

Normal human skin fibroblasts (NHDFs) and patient-derived colon cancer cell line (JC-011) were seeded at 1.0 × 10^4^ cell numbers on collagen-coated 96 well plates and cultured for 1 day at 37 °C and 5 % CO_2_ condition using the 200 µl of Dulbecco modified Eagle’s medium (DMEM: 08458-16, nacalai tesque, Kyoto, Japan) or the 1:1 mixed medium of RPMI1640 (189-02025, WAKO, Osaka, Japan) and Ham’s F-12 (087-08335, WAKO, Osaka, Japan), respectively. Both media were supplemented with 10 % fetal bovine serum (FBS) (10270-106, Thermo Fisher Scientific, MA, USA) and 1 % antibiotics. The 0.2 M K_2_PtCl_4_, NaCl, or glutaraldehyde (020-34385, KISHIDA CHEMICAL, Osaka, Japan) in PBS was diluted to 0.01–5.0 mM by each culture medium. After the medium removing and rinsing the cells with PBS, 200 µl of each fresh medium containing the substances was added, and subsequently incubated for 1 day. After the incubation, the culture media were removed and then the cells were rinsed with 100 µl of fresh media and subsequently the cell viability was estimated by a WST-8 reagent (07553-15, nacalai tesque, Kyoto, Japan). To evaluate effect of the substances on cell proliferation, NHDFs were seeded at 1.0 × 10^4^ cell numbers in 48 well plates (3830-048, IWAKI, Shizuoka, Japan) and incubated in 500 µl of DMEM for 1 day at 37 °C and 5 % CO_2_ condition. After the medium removing and cell rinsing with PBS, 500 µl of fresh DMEM containing substances was added for 1–3 days of culture. The living cell number was estimated by a WST-8 reagent at each time point.

### Healing property of Col-Pt gels

To evaluate self-healing property of Col-Pt gels, 1 ml of 0.5 mM Pt(II)-1.0 wt% CNF gel was prepared in a circular mold using K_2_PtCl_4_. After cutting the gels in half, 20 µl of PBS, 1.0 mM K_2_PtCl_4_ in PBS, or 1.0 wt% CNF solution was added to the cut gel surfaces and then the gels were attached together for 1 day of incubation at room temperature. The photos were taken after lifting the gels.

### Culture of cancer tissue-originated spheroid

The clinical features of the patients from whom human CRC organoids were derived have been described in a previous report (*34*). The organoids were prepared according to the CTOS method (*32*). In brief, resected xenograft tumors were mechanically dissociated, then partially digested with 0.26 U/ml of Liberase DH (Roche, Mannheim, Germany). The organoids were collected using 100-and 40-μm cell strainers (BD Falcon, Franklin Lakes, NJ, USA). The organoids were cultured in StemPro hESC (Invitrogen, Carlsbad, CA, USA).

### Plasmid construction and gene transfer to the organoids

Organoids were transfected with the expression vector pPiggyBac (PB)-Ubc.eGFP-neo and pCMV-hyPBase. Electroporation was performed in 2-mm gap cuvettes at 150 V for 5 msec, using Type II NEPA21 electroporator (Nepa Gene, Chiba,Japan). After transfection, organoids were selected with G-418 (Roche Applied Science) and maintained in the medium containing G-418.

### 3D-culture of organoids in Col-Pt gels

The 50 μl of 0.5 mM Pt(II)-0.2 wt% CNF solution was added in 24 microwells and then incubated at 37 °C for 30 min to prepare the bottom gels. Five of the EGFP-labeled organoids were added to 50 μl of 0.5 mM Pt(II)-0.2 wt% CNF solution and then dropped on the surfaces of the bottom gels surfaces. After incubation at 37 °C for 30 min to fabricate assembled Col-Pt gels containing the organoids, 2 ml of StemPro hESC (Invitrogen, Carlsbad, CA, USA) was added and then incubated it for 7 days. The gels were incubated with 1 μg/mL propidiumiodide (PI) (Molecular Probes) and 2 μg/mL Hoechst33342 (Molecular Probes) at 37 °C for 15 min to visualize the morphology of the organoids and the dead cells. After washing the gels with Hank’s balanced salt solution (HBSS, Merck, Darmstadt, Germany), the morphologies of over 10 organoids were observed by Leica DMi8 microscope (Leica Microsystems, Wetzlar, Germany) and then the organoid shapes were categorized for the following two types.

1. There are clear boundaries, and no boundaries are crossed.
2. The boundaries are indistinct, and spike and cell migration as a cell population toward the outer boundary were observed.

Fluorescence images were obtained using confocal microscopy (TCS SPE; Leica Microsystems, Wetzlar, Germany). The control soft gels without Pt (II) ions were constructed by the commercial 0.1 wt% type I collagen solution by neutralization and heating at 37 °C using a collagen gel culturing kit (Cellmatrix®, Nitta Gelatin Inc., Osaka, Japan). The images of the categories were indicated in Fig. S8.

## S2. Effect of TM species on gelation

**Table S1.**
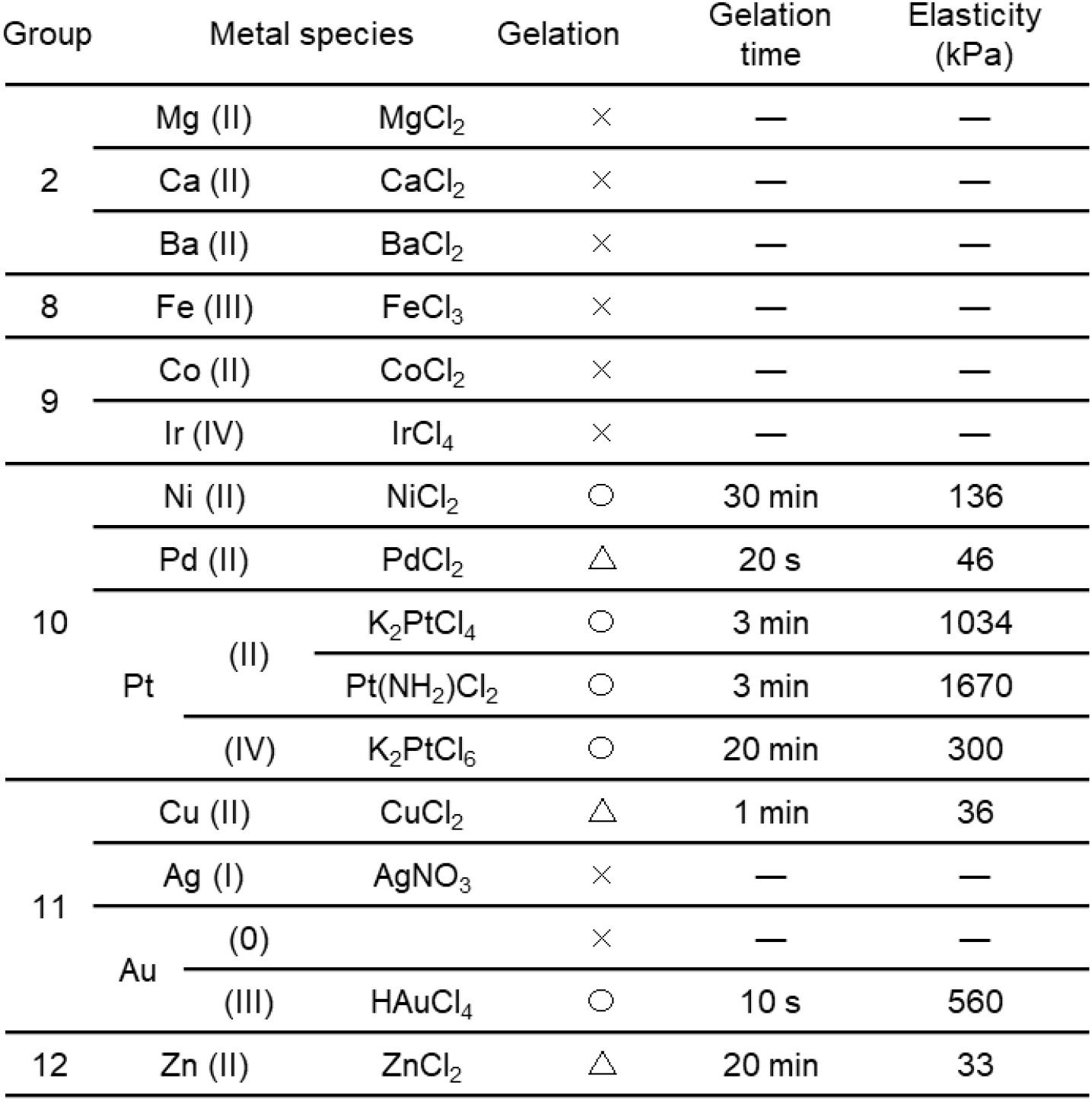
Effect of metal species on gelation property of Col-TM gels fabricated by mixing 0.5 wt% CNF and 0.5 mM TM ions.

## S3. CD spectrum of Col-Au gels

**Fig. S1.**
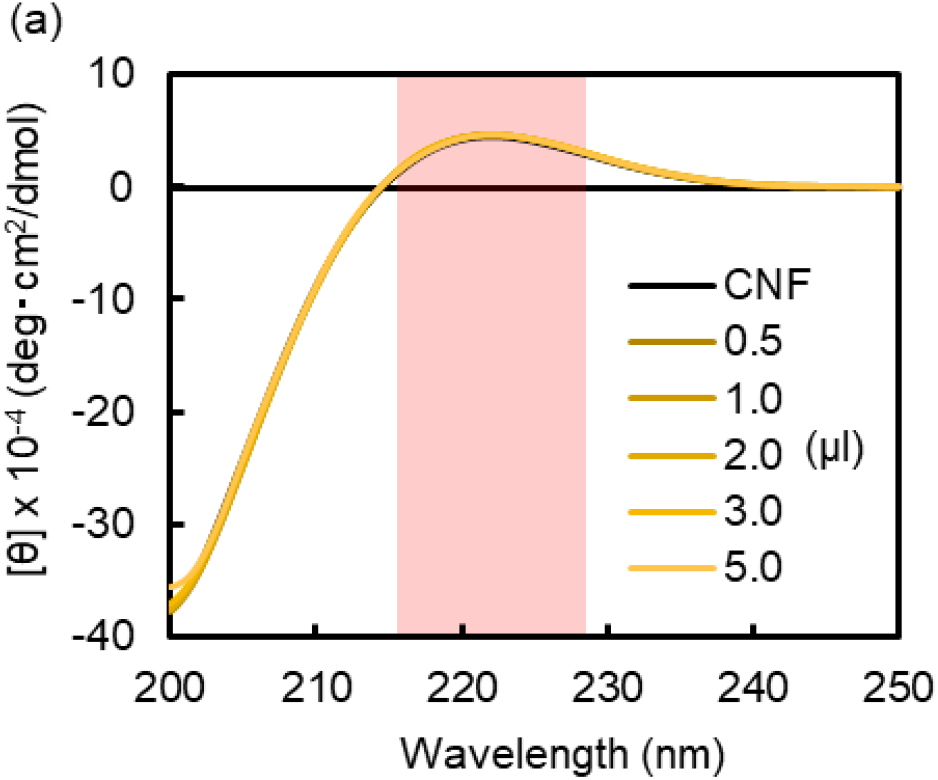
CD spectra of CNF solution with different concentration of HAuCl_4_ at r.t.

## S4. Effect of collagen species and type on gelation

**Table S2.** Effect of collagen species and type on gelation property of Col-Pt gels.

| Species | Type | Triple helix | Gelation |
| --- | --- | --- | --- |
| Pig | Col I | ○ | ○ |
|  | Col III | ○ | ○ |
|  | Tropo Col I | ○ | ○ |
| Chicken | Col II | ○ | ○ |
| Bovine | Col I | ○ | ○ |
| Human | Col I | ○ | ○ |
|  | Col IV | ○ | ○ |

## S5. Effect of polymer type on gelation

**Table S3.** Effect of polymer type on gelation property of Col-Pt gels.

| Polymer | M<sub>W</sub> | Helix | Gelation |
| --- | --- | --- | --- |
| Gelatin | 100,000 | × | × |
| (POG)<sub>10</sub> | 2,700 | ○ | × |
| Laminin | 865,000 | × | × |
| Fibronectin (0.1 wt%) | 250,000 | × | × |
| Heparin | 12,000 | × | × |
| Alginate | — | × | × |
| Polyacrylate | 250,000 | × | × |
| DNA | — | ○ | × |

## S6. Effect of gelatin concentration on gelation

**Table S4.** Effect of gelatin concentration on gelation property of Col-Pt gels.

| Gelatin concentration<br>(wt%) | Pt(II) concentration<br>(mM) | Gelation |
| --- | --- | --- |
| 5.0 | 1.0 | × |
|  | 5.0 | × |
|  | 10 | × |
|  | 50 | × |
| 10 | 1.0 | × |
|  | 5.0 | △ |
|  | 10 | ○ |
| 20 | 1.0 | × |
|  | 5.0 | ○ |
|  | 10 | ○ |

## S7. Viscoelasticity of CNF and gelatin solutions

**Table S5.** Viscoelasticity of CNF and gelatin solutions.

| Substance | wt% | Viscoelasticity<br>(mPa·S) |
| --- | --- | --- |
| CNF | 0.2 | 22 |
|  | 0.5 | 121 |
|  | 1.0 | 429 |
| Gelatin | 5.0 | 1.6 |
|  | 10 | 5.0 |
|  | 20 | 22 |

## S8. Pt L_3_-edge X-ray absorption spectroscopy of Col-TM gels

**Fig. S2.**
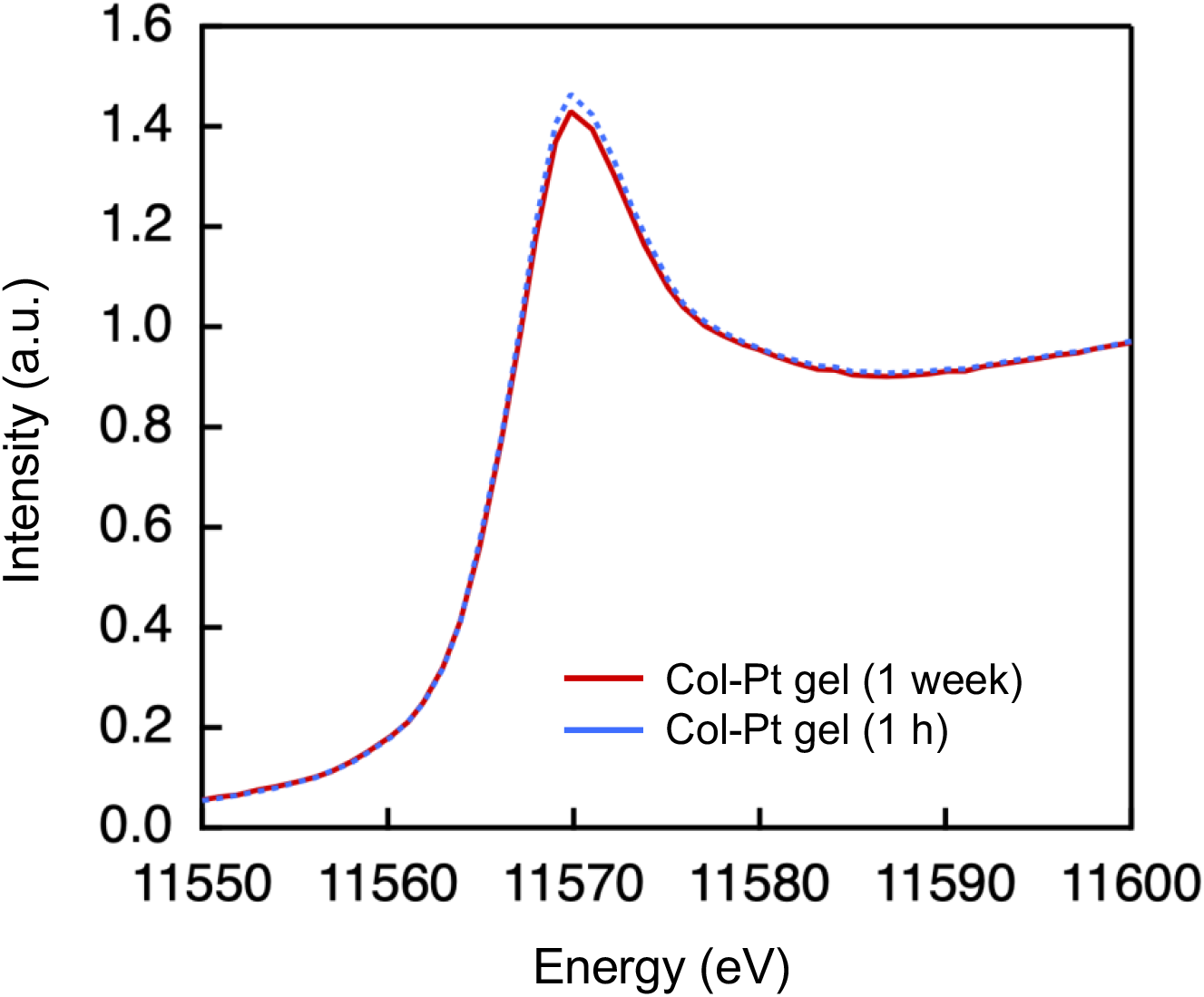
XANES spectra of Col-Pt gels after incubation for 1 h (blue dotted line) and 1 week (red line) at 4 °C

**Fig. S3.**
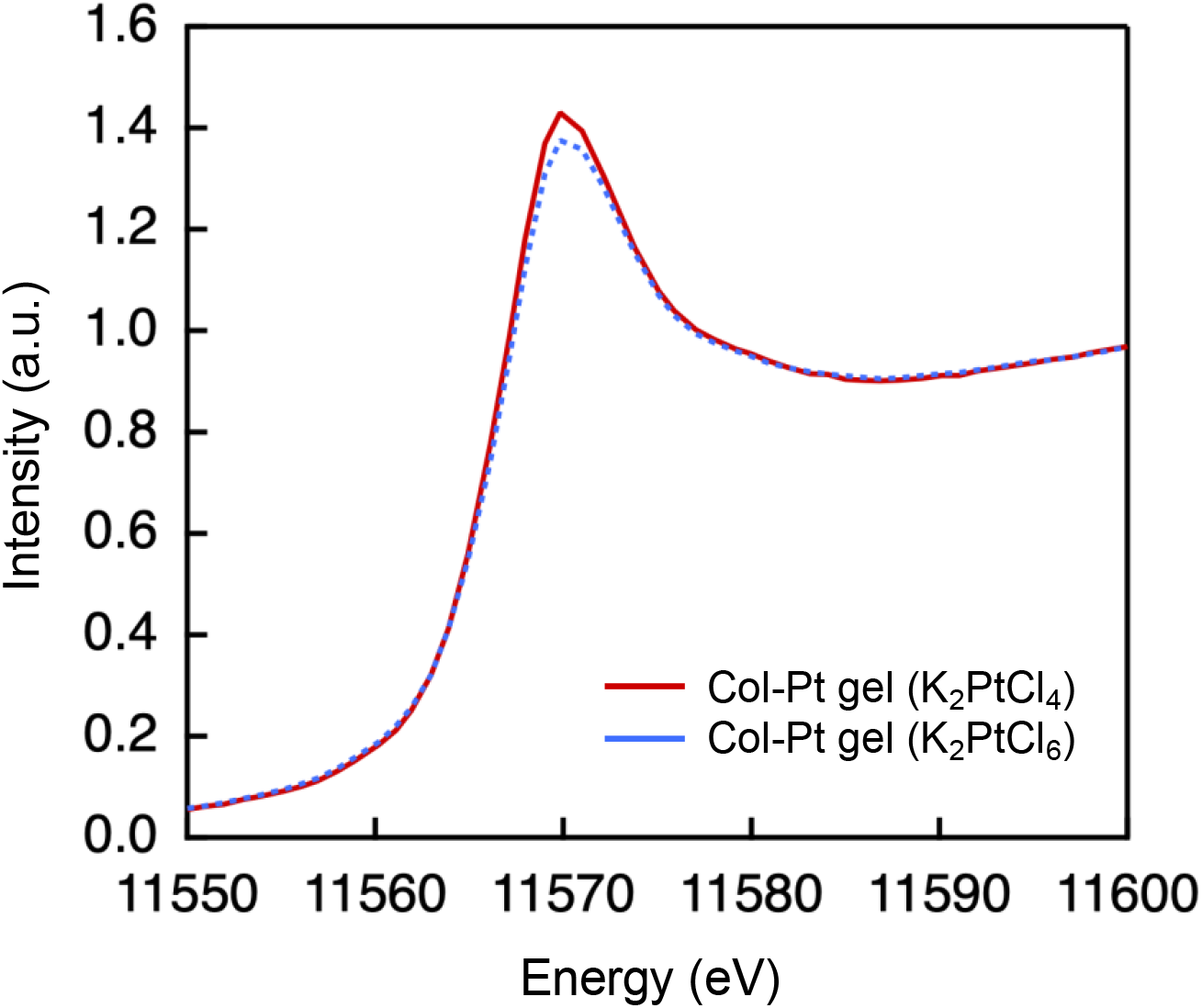
XANES spectra of Col-Pt gels prepared from K_2_PtCl_4_ (red line) and K_2_PtCl_6_ (blue dotted line).

**Fig. S4.**
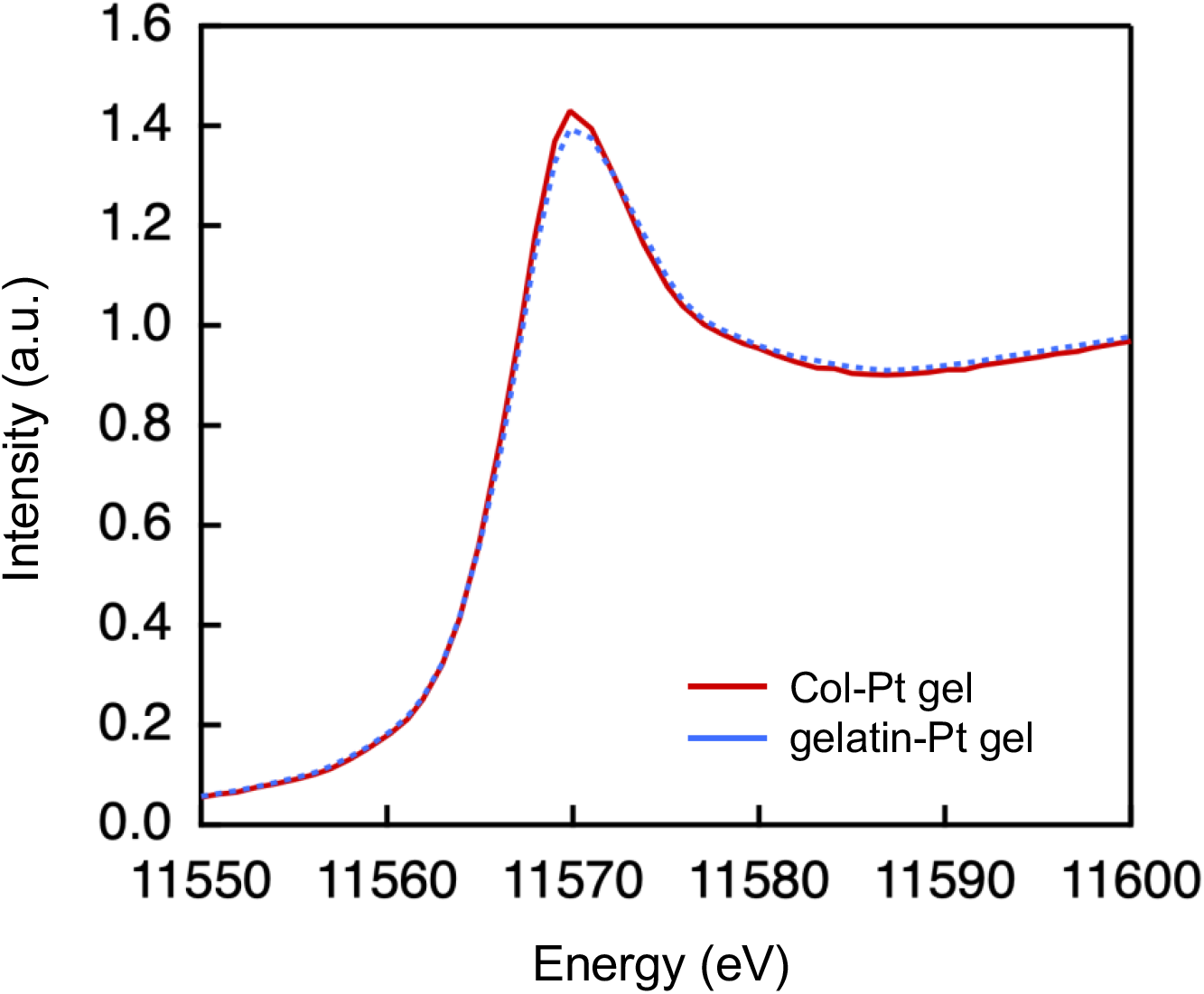
XANES spectra of Col-Pt gel (red line) and gelatin-Pt gel (blue dotted line).

**Fig. S5.**
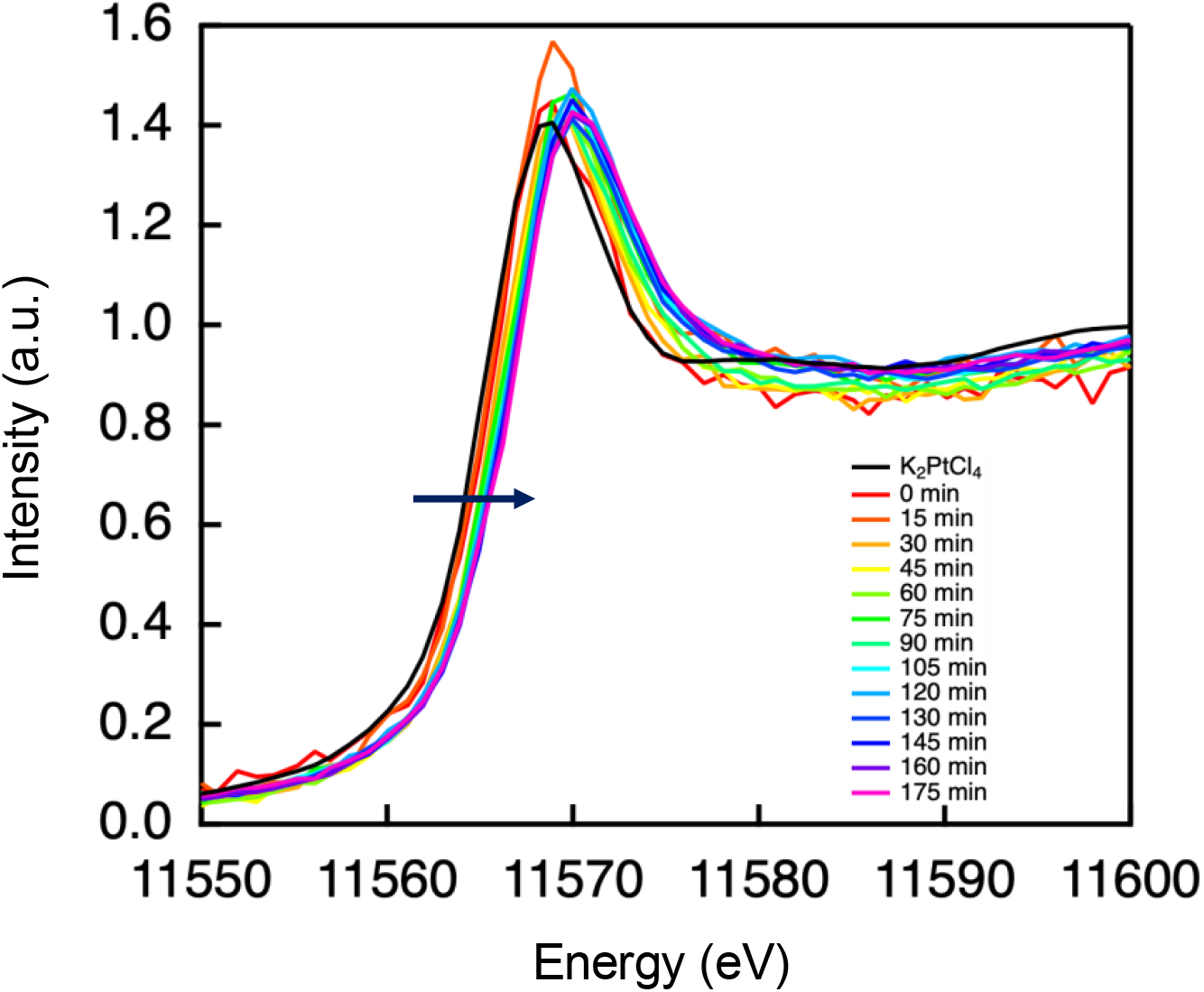
Time-course of XANES spectra of (POG)_10_ + K_2_Pt(II)Cl_4_ in PBS. The times shown inset signify the start time of the measurements. XAS measurement takes approximately 15 min for 1 scan.

## S9. ICP data of Col-Pt gels

**Fig. S6.**
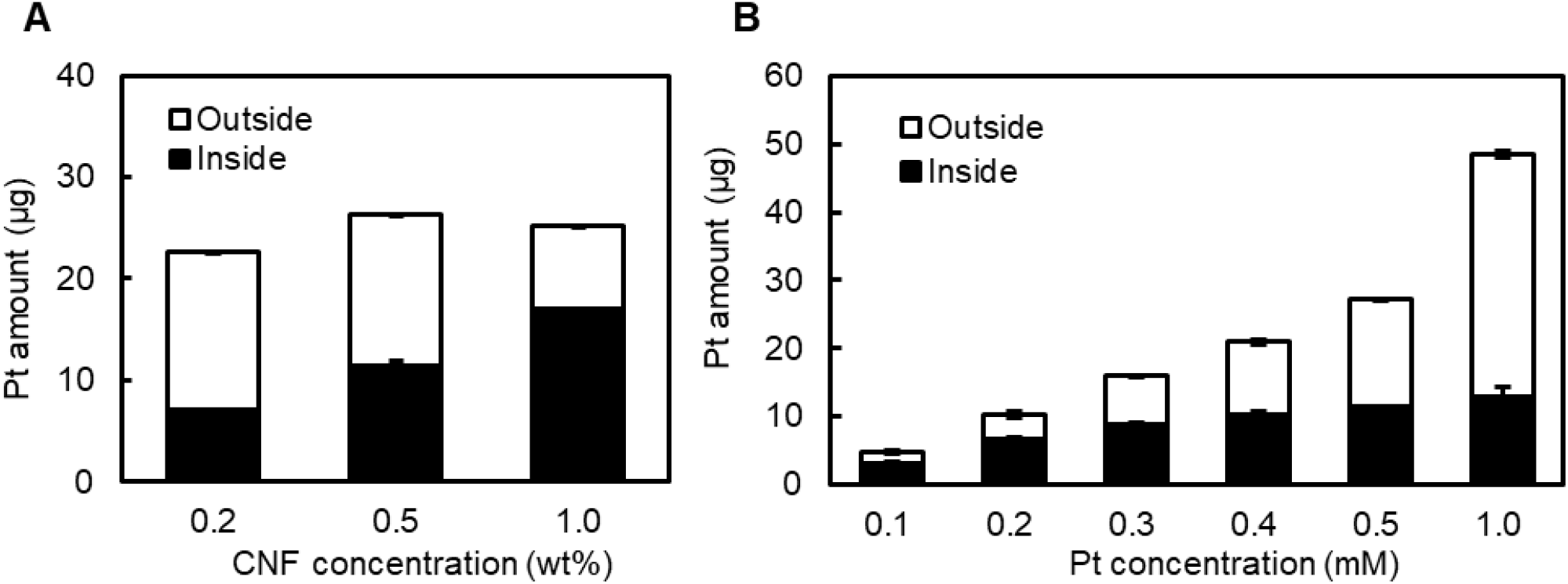
Estimation of the amount of Pt(II) ions inside and outside Col-TM gels (n=3). Col-TM gels were prepared by (**A**) 0.5 mM K_2_Pt(II)Cl_4_ and 0.2–1.0 wt% CNF solutions and (**B**) 0.5 wt% CNF solution and 0.1–1.0 mM K_2_Pt(II)Cl_4_. The obtained gels were washed with 1 ml of PBS for 3 times and then measured the amount of Pt(II) ions inside the gels (inside) and whole washing PBS (outside).

## S10. Cell viability assay

**Fig. S7.**
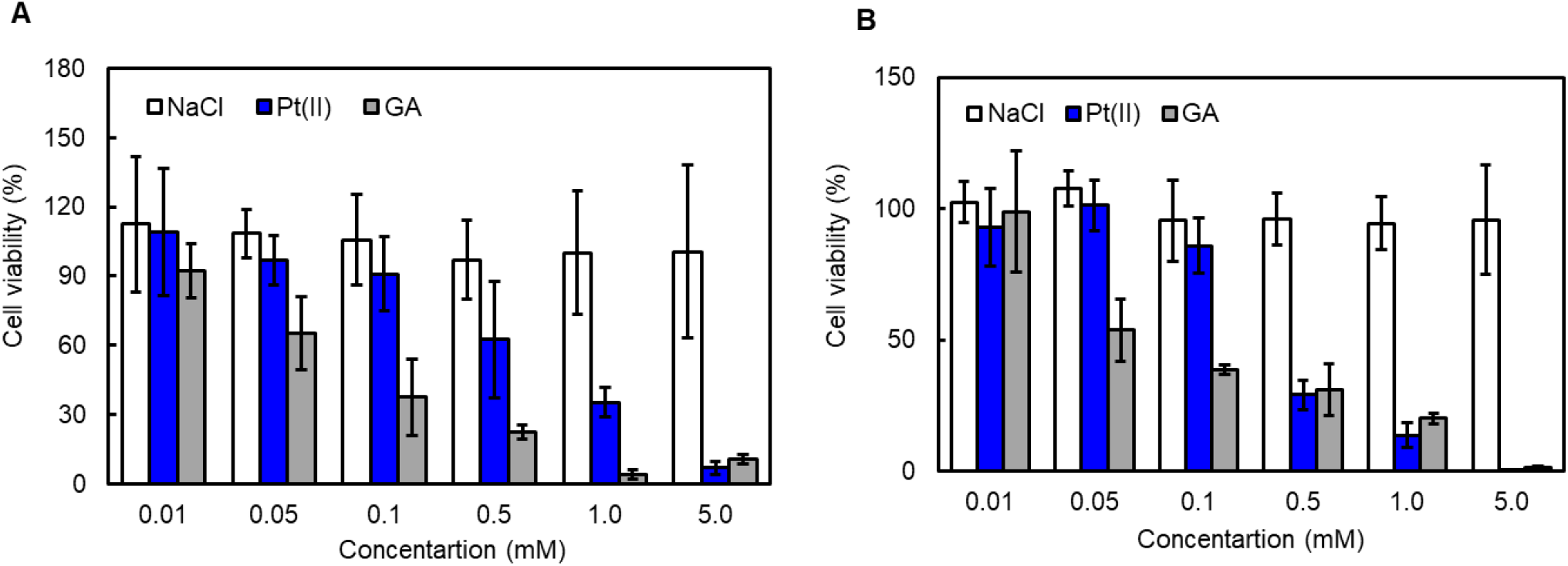
Cell viability of (**A**) JC-011 and (**B**) NHDFs in DMEM containing 10% FBS and NaCl, glutaraldehyde (GA) or K_2_Pt(II)Cl_4_ for 3 days of culture without medium change (n=3).

## S11. 3D culture of organoids in Col-Pt gels

**Fig. S8.**
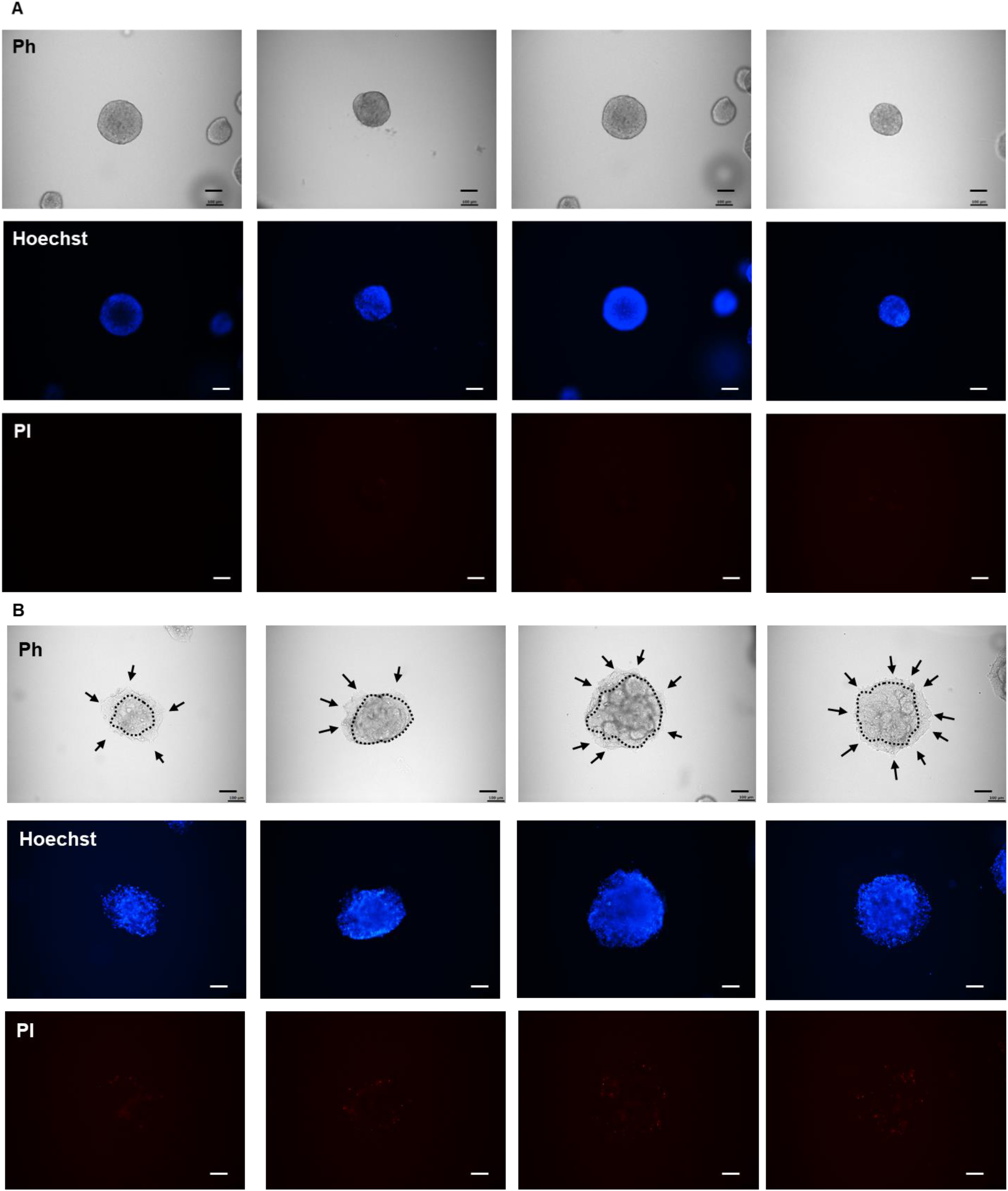
Phase contrast, Hoechst and PI images of CTOS cultured in (**A**) control soft collagen gels and (**B**) stiff Col-Pt gels constructed by 0.2 wt% CNF with 0.5 mM K_2_PtCl_4_ solutions. Black arrows and dashed lines indicate spike/cell migrations and boundary, respectively. Scale bars are 100 µm.

**Fig. S9.**
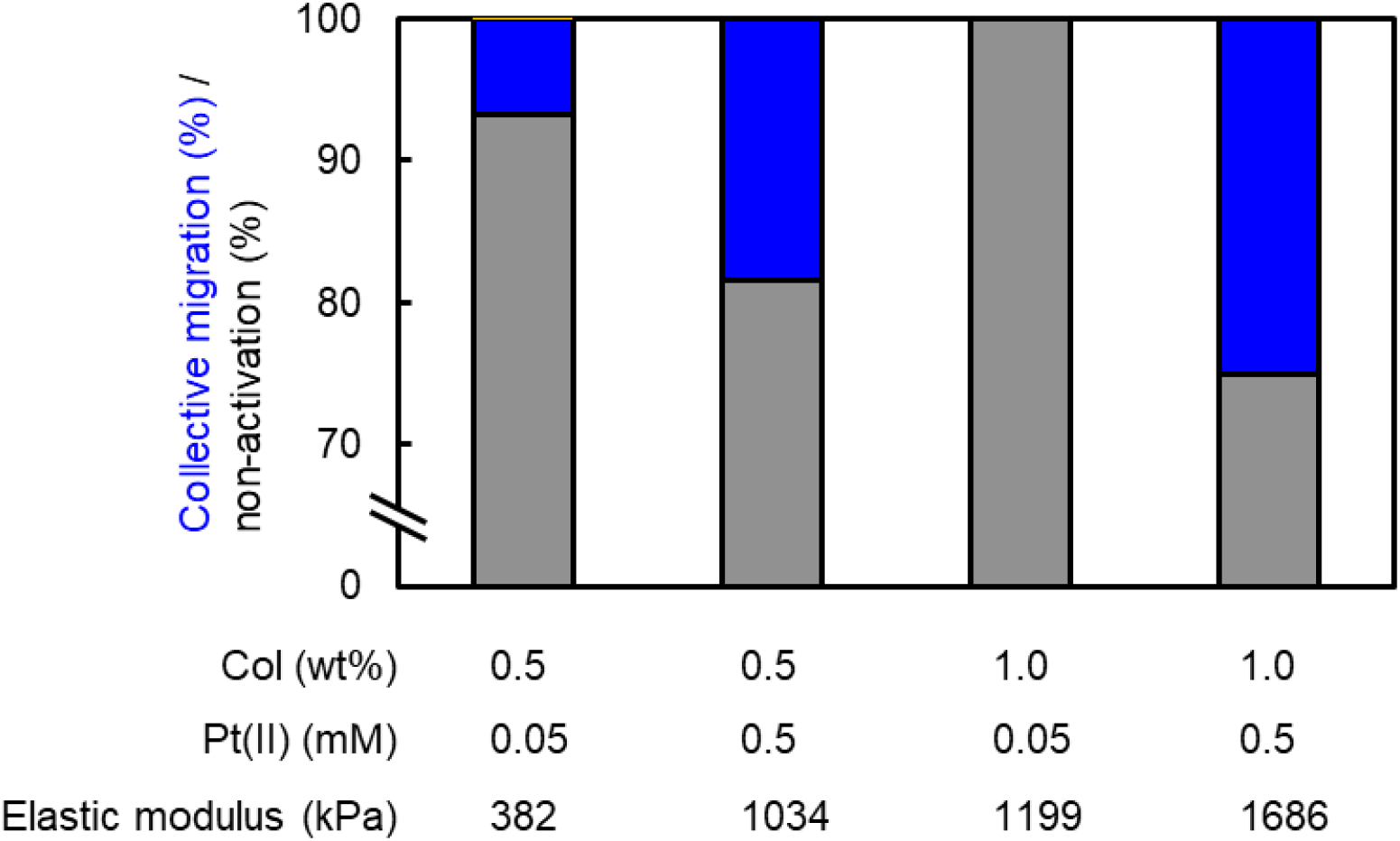
Mean percentage of both collective migration and non-activation of organoids after 3D culture inside the gels constructed by varied Col and Pt(II) ion concentrations (n=5∼6).

## S12. Gelation videos of Col-TM gels

**Movie S1.**
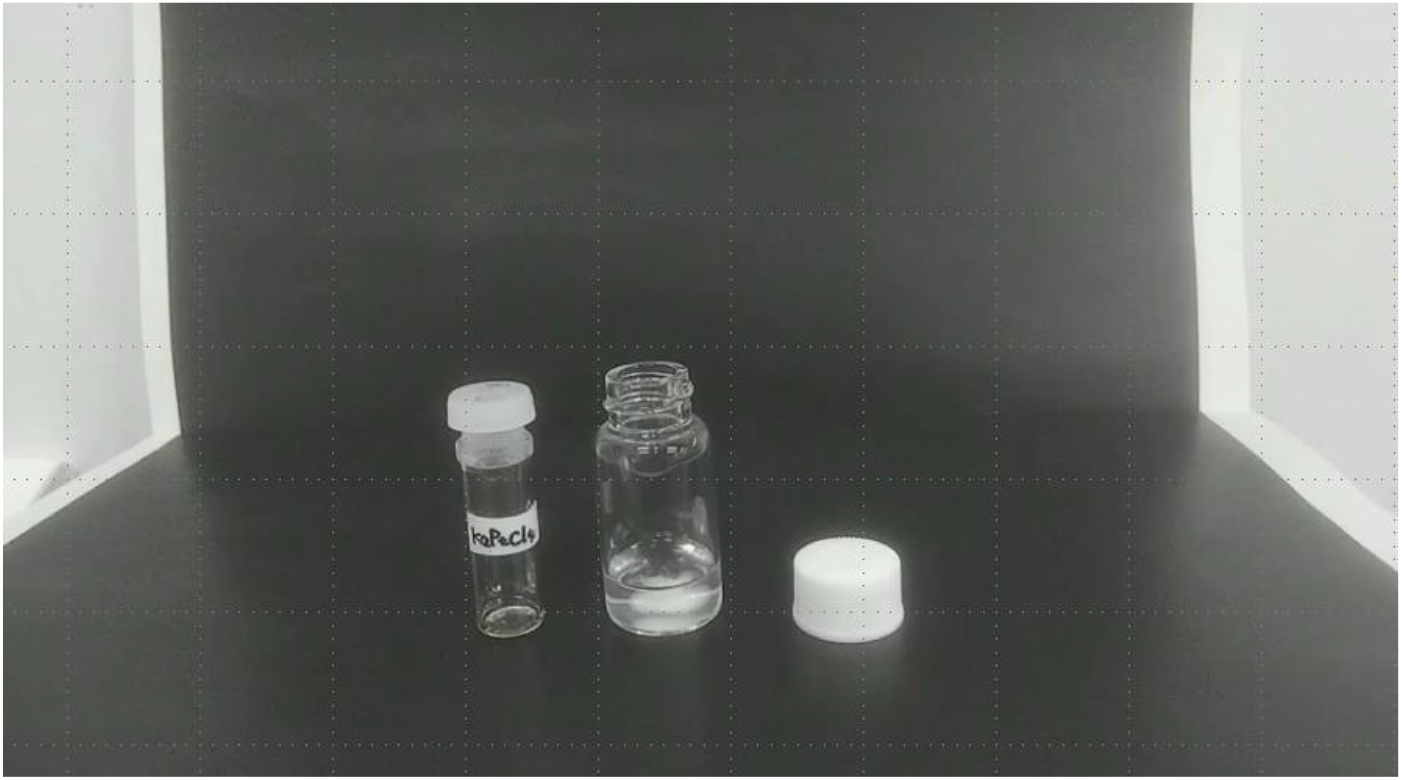
Gelation video of 0.5 wt% CNF solution by mixing with 0.5 mM K_2_Pt(II)Cl_4_ at r.t. Speed: x2.

**Movie S2.**
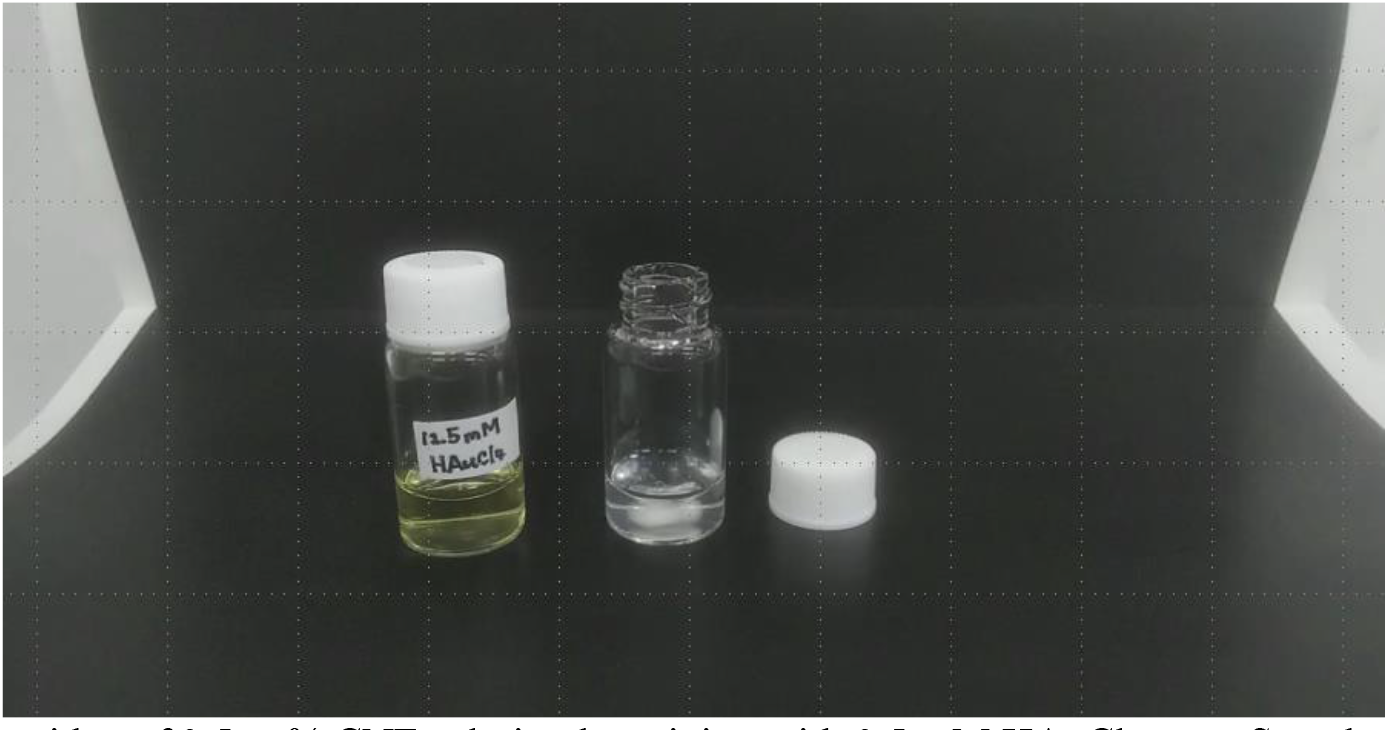
Gelation video of 0.5 wt% CNF solution by mixing with 0.5 mM HAuCl_4_ at r.t. Speed: x2.

**Movie S3.**
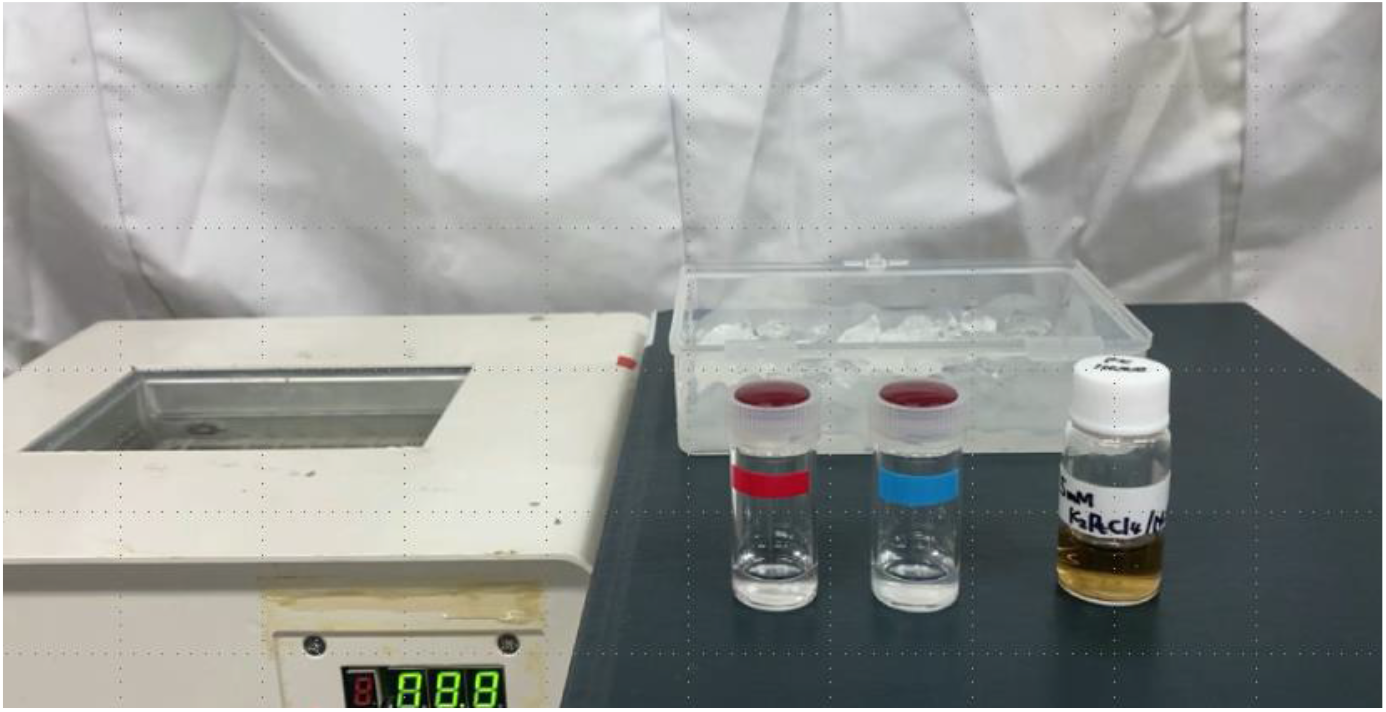
Gelation comparison between 0.5 wt% CNF (blue vial) and 0.5 wt% gelatin (red vial) solutions by mixing with 0.5 mM K_2_Pt(II)Cl_4_ at r.t. Speed: x2.

